# Metabolic maintenance of breast cancer cells and metastases through E-cadherin/YAP–dependent pyruvate carboxylase expression

**DOI:** 10.64898/2026.04.13.718309

**Authors:** Kuppusamy Balamurugan, Jonathan M. Weiss, Lois McKennett, Shikha Sharan, Brad A. Gouker, Donna O. Butcher, David A. Scheiblin, Elijah F. Edmondson, Duncan Donohue, Stephen J. Lockett, Laura Bassel, Daniel W. McVicar, Esta Sterneck

## Abstract

Epithelial-mesenchymal transition (EMT) and glycolytic metabolism are well-characterized drivers of cancer progression and metastasis. However, most primary breast tumors and metastases express E-cadherin and the epithelial phenotype is associated with mitochondrial oxidative metabolism, yet the causality and relevance of these relationships and their underlying mechanisms remain poorly understood. Using a 3D culture model with mechano-stimulation, we found that E-cadherin promotes mitochondrial oxidative phosphorylation (OXPHOS) while reducing oxidative stress. Through pharmacological and genetic manipulations of inflammatory breast cancer (IBC) and/or triple negative breast cancer (TNBC) cell lines, we identified pyruvate carboxylase (PC) as an E-cadherin effector. Critically, restoring PC in E-cadherin-silenced cells rescued mitochondrial oxygen consumption and protection from oxidative stress. Co-expression of E-cadherin and PC was confirmed in breast cancer tissues and experimental lung metastases. Mechanistically, E-cadherin induced PC expression and OXPHOS via AKT-mediated activation of YAP/ /TEAD transcription factors, which are better known as supporting EMT. Clinically relevant AKT and TEAD inhibitors reduced both PC expression and oxidative respiration. Importantly, PC inhibition as monotherapy attenuated established experimental lung metastases and primary tumor burden in mice. Taken together, these findings reveal that E-cadherin-mediated cell-cell adhesions directly support mitochondrial metabolism through AKT–YAP/TEAD–PC signaling, identifying a therapeutic vulnerability in metastatic epithelial TNBC.

## Introduction

Epithelial-mesenchymal transition (EMT) is a well-characterized mechanism of cancer progression and metastasis, in which epithelial markers such as E-cadherin are downregulated while mesenchymal markers including vimentin are induced. EMT is regulated by a set of transcription factors and has been linked to increased stemness and invasiveness (1, 2). Accordingly, EMT gene signatures are enriched in metastatic tumors (3) and have been implicated in multiple steps of the metastatic cascade (4, 5). However, we previously demonstrated that E-cadherin mRNA abundance does not reliably predict E-cadherin protein expression in cancer cells (6). These findings align with a growing body of evidence indicating that cancer cell dissemination does not require a complete EMT but rather involves epithelial-mesenchymal (E/M) plasticity or hybrid states, which are also associated with cancer stem cell (CSC) properties and circulating tumor cell (CTC) survival (1, 7, 8).

In breast cancer (BC), most invasive ductal carcinomas and metastases retain E-cadherin protein expression, including the highly aggressive inflammatory breast cancer (IBC) subtype (9–11). Multiple studies have reported a positive correlation between E-cadherin expression and metastasis (9, 12–14), and that E-cadherin can contribute to collective cell migration and metastasis establishment, chemotherapy resistance, and cancer survival in CTC clusters (6, 15–17). Additionally, the epithelial phenotype has been linked to protection from oxidative stress, at least in part through engagement of the serine synthesis pathway (16, 18). Nonetheless, compared to the EMT process, the causal relationships and molecular mechanisms through which E-cadherin contributes to breast cancer aggressiveness and metastasis remain incompletely defined.

Beyond EMT, metabolic plasticity has emerged as an additional hallmark of cancer progression (8, 19). Although aerobic glycolysis (the Warburg effect) in solid tumors has been linked to EMT induction, immune evasion, and tumor progression (20), mitochondrial metabolism, including the tricarboxylic acid (TCA) cycle and oxidative phosphorylation (OXPHOS), is increasingly recognized as contributing to chemoresistance and metastatic competence (8, 21, 22). Epithelial cancer cells have been reported to display enhanced oxidative metabolism (18, 23), but the mechanistically causal nature of this relationship is poorly understood.

In this study, we investigated the potential causal link between E-cadherin expression and mitochondrial metabolism using the 3D “emboli” culture (EmC) protocol. This paradigm employs slightly viscous media and gentle orbital rocking in ultra-low attachment plates to promote stable cell-cell adhesions, resulting in compact multicellular aggregates which recapitulate select ultrastructural features of nuclei, mitochondria and lipid droplets as seen *in vivo* (24). Using this approach, we previously showed that basal epithelial breast cancer cell lines, i.e. the triple negative BC (TNBC) MDA-MB-468 and SUM149 as well as HER2+ IBC3 cells (the latter two also representing IBC), exhibit increased OXPHOS when cultured in 3D compared with 2D cultures, whereas mesenchymal TNBC cell lines showed reduced OXPHOS (24). Here, we identify an E-cadherin dependent signaling pathway that induces pyruvate carboxylase (PC) expression via the YAP/TEAD transcription factors. Consistent with a functional role of this pathway, PC inhibition reduced experimental lung metastases in a preclinical mouse model. Accordingly, E-cadherin and PC were co-expressed in vivo in both experimental lung metastases and breast cancer specimens. Together, these findings support a role for E-cadherin-PC signaling in metastatic progression and support PC and glutamine pathway as a therapeutically actionable metabolic vulnerability in epithelial breast cancer.

## RESULTS

### E-cadherin promotes mitochondrial respiration in breast cancer cells

To address the potential role of E-cadherin in the adaptation of mitochondrial metabolism, we assessed the effect of E-cadherin depletion through stable expression of shRNA targeting its mRNA *CDH1* (**Figure 1A**). As shown in **Figure 1B**, silencing of *CDH1*/E-cadherin expression did not perturb emboli formation by SUM149 cells, while smaller aggregates were formed by E-cadherin-depleted MDA-MB-468 and IBC-3 cells, which was also reflected in fewer cells recovered from emboli after 3 days of culture (**Figure 1C**). These data indicate various degrees of dependence on E-cadherin for emboli formation and/or survival under these conditions, with SUM149 cells exhibiting least dependence on E-cadherin for cell-cell adhesion. Unless indicated otherwise, all data presented here are from cells cultured for 3-5 days as emboli/aggregates in EmC and which were selectively isolated by brief low-speed centrifugations (24). Next, cells from these emboli and aggregates were dissociated to assess their oxidative consumption rate (OCR) as a measure of OXPHOS. Cells with reduced E-cadherin expression exhibited reduced OCR (**Figure 1D**) according to the degree of shRNA silencing efficiency by each of the two shRNAs (shRNA-2 was more efficient than shRNA-1), and between cell lines (**Figure 1A**). The extracellular acidification rate (ECAR) as a measure of glycolytic activity was induced (SUM149, MB468) or unchanged (IBC-3) by E-cadherin depletion (**Figure 1E**). Combining data from three independent experiments, we calculated OCR/ECAR ratios which were similarly compromised in all three cell lines upon E-cadherin knockdown (**Figure 1F**). In addition, E-cadherin depleted cells showed more DCF fluorescence (**Figure 1G**), a measure of reactive oxygen species (ROS), which is in agreement with the reported role of E-cadherin in ROS protection (16) and of efficient oxidative phosphorylation reducing ROS (25–27). Because only cells from aggregates were analyzed and E-cadherin depleted SUM149 cells continued to form large emboli in EmC, these data indicate a direct role of E-cadherin in mitochondrial metabolism.

**Figure 1.**
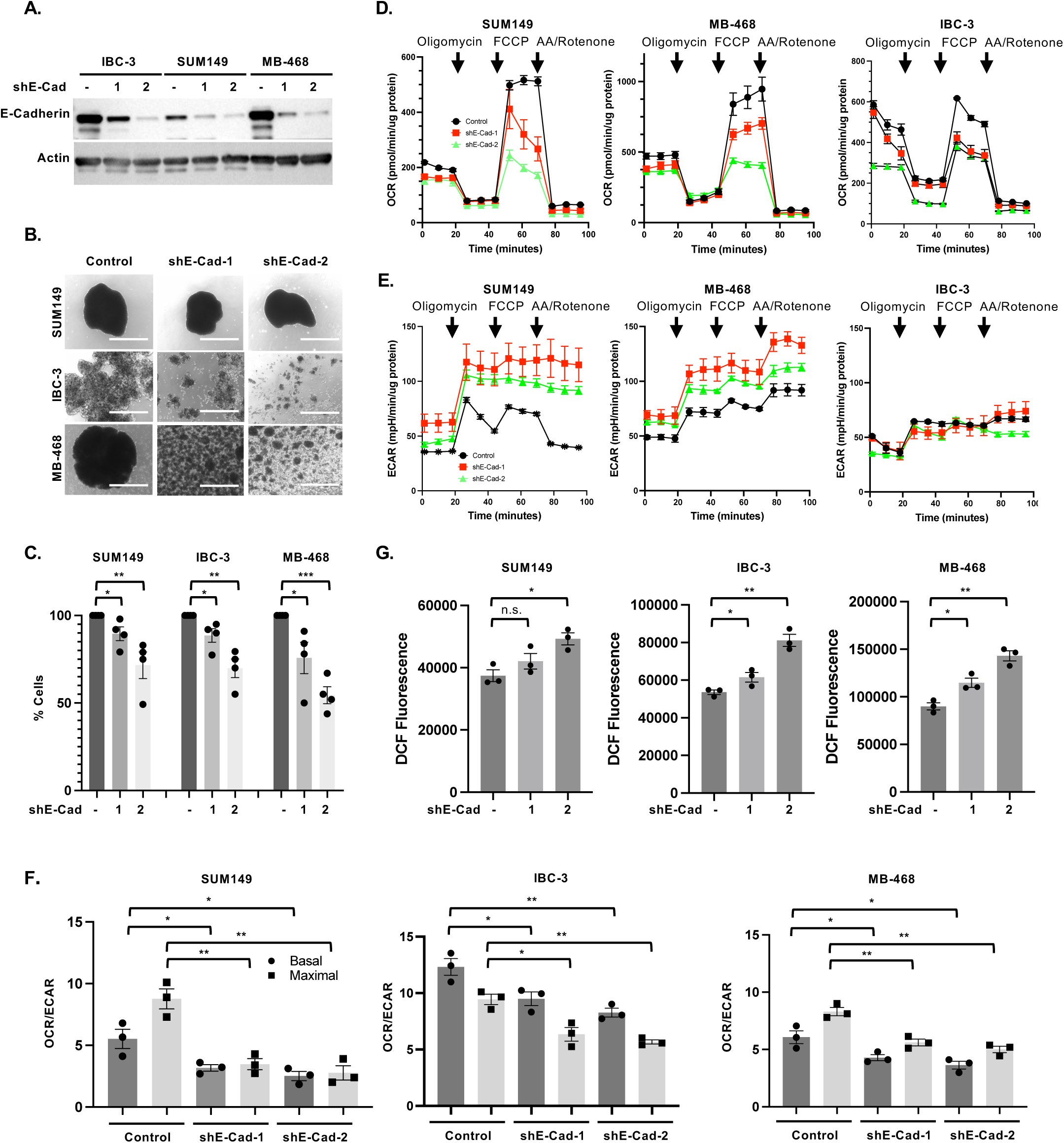
E-Cadherin promotes mitochondrial respiration in breast cancer cells. **(A)**. Western analysis showing E-Cadherin levels in the indicated pools of cells with stable expression of one of two independent *CDH1* targeting shRNA (shE-Cad #1 and #2) in comparison to control, all cultured in 2D. **(B)** Bright-field images of SUM149, MB-468 and IBC-3 cell as in panel A after 3 days in EmC (scale bar =1 mm). **(C)** Quantification of viable cells associated with emboli/cell aggregates shown in panel B, expressed as percent viable cells by E-cadherin-depletion compared to control cells (n = 3). **(D)** Representative Seahorse analysis of the oxygen consumption rate (OCR) over time by cells from cultures as in panel B. **(E)** Representative Seahorse analysis of the extracellular acidification rate (ECAR) profiles from the experiment shown in panel D. **(F)** Basal and maximal OCR/ECAR ratios of the indicated cell lines derived from data as in panels D and E (n=3 independent experiments). **(G)** Fluorometric determination (arbitrary units) of ROS in the indicated cell lines after 3 days of culture in EmC (n=3). Bar graphs are mean ± SEM. \**P*<0.05, \*\**P*<0.01, \*\*\**P*<0.001; n.s., not significant.

### E-cadherin promotes expression of the mitochondrial pyruvate carboxylase

To address the mechanism by which E-Cadherin promotes OXPHOS, we interrogated proteomic data from our prior characterization of cells in EmC (24) which suggested increased levels of pyruvate carboxylase (PC), an anaplerotic enzyme that converts pyruvate into oxaloacetate. This enzymatic activity feeds the TCA cycle, promotes OXPHOS, and indirectly supports antioxidant pathways (28–32). Therefore, we hypothesized that E-Cadherin may promote OXPHOS through PC. First, we validated that all three cell lines increased PC expression in EmC compared to cells in 2D culture, both, at the level of mRNA and protein (**Figures 2A-B**). Compared to control cells, E-Cadherin knockdown significantly compromised PC mRNA and protein levels in all three cell lines in EmC (**Figures 2C-D**). Transient ectopic expression of E-cadherin in the mesenchymal TNBC cell lines MDA-MB-231-LM2 and SUM159 did not significantly change the appearance of cell aggregates in EmC but increased the number of viable cells in MB-231-LM2 cells after 3 days of culture (**Figure S1A**) and was sufficient to induce expression of PC (**Figure 2E**). These data demonstrate a role of E-cadherin in supporting *PC* expression. While E-cadherin is abundantly expressed in normal mammary epithelial cells (33), PC expression is generally absent but induced in breast cancer (31, 34). To determine the relationship of PC and E-Cadherin *in vivo*, we performed immunofluorescence staining of SUM149 experimental lung metastases (**Figure 2F**). Quantification of co-expression in four independent lesions (**Supplemental Figure S1B**) showed that the great majority of PC expressing cells also expressed E-cadherin (**Figure 2G**). These data demonstrate that the epithelial phenotypes favor activation of the *PC* gene in cancer cells *in vivo*. Next, we extended this analysis to human breast cancer specimens and found that PC expression was enriched in epithelial E-cadherin expressing cells compared to mesenchymal cells and that such co-expression was seen in all the three subtypes. (TNBC, HER2+ and ER+) (**Figures 2H-I** and **Supplemental File 1**). To determine if co-expressing cells are potentially metastatic, we analyzed circulating tumor cells (CTCs) in mice with SUM149 xenograft tumors and observed that some CTCs co-expressed both proteins (**Figure 2J**), demonstrating that such cells can leave the primary tumor. Taken together, these data demonstrate preferential co-expression of PC with E-cadherin in vivo.

**Figure 2.**
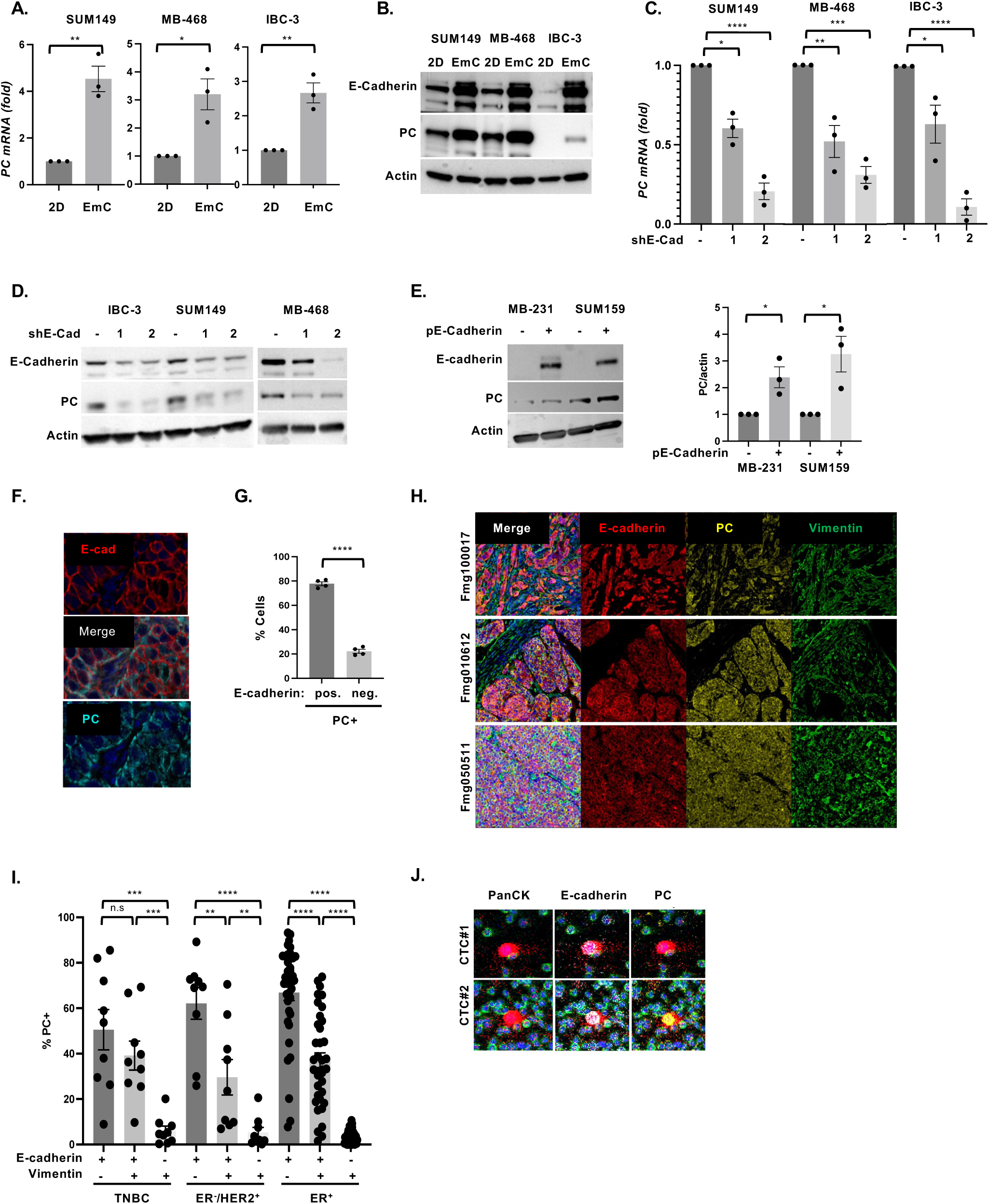
E-cadherin promotes expression of the mitochondrial pyruvate carboxylase. **(A)** qRT-PCR analysis of PC expression in SUM149, MB-468 and IBC-3 cells after 3 days of culture in 2D or EmC. **(B)** Representative Western analysis of the indicated proteins in cells as in panel A. Actin served as a loading control. **(C)** qRT-PCR analysis of PC expression in SUM149, MB-468 and IBC-3 cell lines with stable expression of shE-Cad-1 or shE-Cad-2 compared to controls. **(D)** Western analysis of the indicated proteins in cells as in panel C. **(E)** Representative Western analysis (left panel) of E-cadherin and PC expression in MDA-MB-231-LM2 and SUM159 cells with and without transfection of an E-cadherin expression construct along with quantification of PC protein (right panel) from three independent experiments. **(F)** Immunofluorescent staining of E-Cadherin and PC in a SUM149 experimental lung metastasis. **(G)** Percentages of PC-expressing cells that were positive respectively negative for E-cadherin in four regions representing three SUM149 experimental metastases (35,063 cells total). **(H)** Examples of three TNBC specimens stained with Opal^TM^ multiplex for the indicated proteins visualized by PhenoImager system. **(I)** Percentage of PC-expressing cells (PC+) in breast cancer tissues of the indicated subtypes expressing E-cadherin and/or vimentin (TNBC, n=9, 36,562 cells total; ER^-^/HER2^+^, n=9, 22,982 cells total; ER+, n=37, 102,979 cells total). **(J)** Two examples of circulating tumor cells from peripheral blood of a mouse with SUM149 xenograft tumor analyzed for expression of the indicated proteins by imaging mass cytometry (red, Pan-keratin; white, E-cadherin; yellow, PC; green, CD45 marking murine immune cells). Bar graphs are mean ± SEM; \**P*<0.05, \*\**P*<0.01, \*\*\**P*<0.001, \*\*\*\**P*<0.0001; n.s., not significant.

### Pyruvate carboxylase is responsible for E-cadherin-mediated mitochondrial respiration

To assess the role of PC in BC cell metabolism we utilized the inhibitor ZY-444, which had been shown to bind PC directly and inhibit mitochondrial respiration (29, 32). When added on day two of EmC, after the initial establishment of emboli, the number of viable cells recovered three days later was significantly reduced in all the three BC cell lines (**Figures 3A** and **S2A-B**), which is consistent with the dependence of cells in EmC on OXPHOS (24). Quantitative analysis was warranted because, as documented before (35), the appearance of 3D structures does not always reflect changes in cell viability. For metabolic studies, ZY-444 was added for only 12 hours to 2.5-day old emboli cultures, which had little to no effect on the recovery of viable cells (**Supplemental Figure S2C**). This approach confirmed that ZY-444 reduced the oxidative consumption rate of cells within the 3D structures (**Figures 3B-C** and **Supplemental Figure S2D**) while glycolytic capacity was either increased (SUM149) or unchanged (MB-468) due to ZY-444 (**Supplemental Figure S2E**). In contrast to the epithelial cell lines used here, the mesenchymal TNBC cell lines MDA-MB-231 and SUM159 do not engage OXPHOS when cultured in EmC (24). Upon transient ectopic expression of E-cadherin, however, their oxidative consumption rates were markedly increased and highly sensitive to ZY-444 (**Figures 3D-E**), consistent with the increased PC expression in these cells (see Figure 2D). Finally, we asked whether PC overexpression could rescue the phenotypes of E-Cadherin-depleted cells. As shown in **Figure 3F**, transient PC overexpression increased the number of E-cadherin knockdown cells recovered from EmC after 3 days in culture. Interestingly, ectopic PC modestly increased the levels of E-cadherin protein in these cells (**Figure 3G**), which may be due to feedback regulation or preferential survival/proliferation of E-cadherin-expressing cells. In addition, PC overexpression rescued not only the OCR/ECAR ratios in E-cadherin depleted cells (**Figure 3H)** but also reduced ROS levels (**Figure** 3**I**), consistent with protection from oxidative stress by indirectly supporting the glutathione (GSH) antioxidant system (36). Taken together, these data demonstrate that E-cadherin supports OXPHOS and reduces oxidative stress at least in part through upregulation of PC expression.

**Figure 3.**
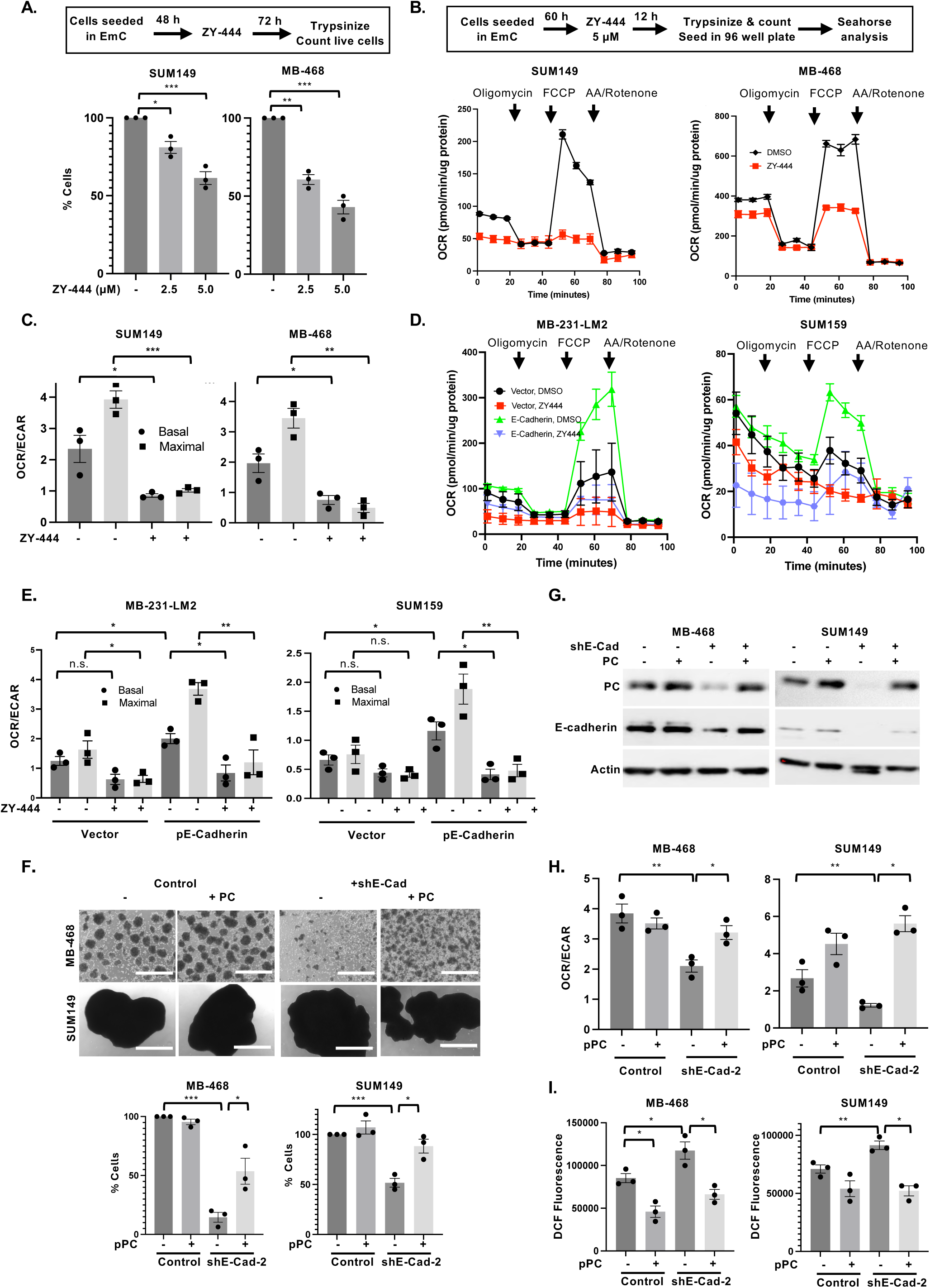
Pyruvate carboxylase is responsible for E-cadherin-mediated mitochondrial respiration. **(A)** Schematic representation of the experimental setup and quantification of % viable cells retrieved from emboli and aggregates compared to vehicle-treated controls (n=3). **(B)** Schematic of experimental setup and representative Seahorse analysis of the oxygen consumption rate (OCR) over time by the indicated cell lines after treatment with ZY-444 (5 µM) for 12 hours. **(C)** Basal and maximal OCR/ECAR ratios of cells as in panel B (n=3). **(D)** Representative Seahorse analysis of the oxygen consumption rate (OCR) over time by the indicated cell lines with or without E-cadherin overexpression and after treatment with ZY-444 as shown in panel B. **(E)** Basal and maximal OCR/ECAR ratios of n=3 independent experiments as shown in panel D. **(F)** Light microscopy images of representative cultures of the indicated cell lines with stable expression of shE-Cad-2 and/or transient overexpression of PC and respective controls (scale bar =1 mm) along with quantification of viable cells after 3 days of culture (n=3). **(G)** Western analysis of the indicated proteins in cells as shown in panel F. **(H)** Basal OCR/ECAR ratios of cells from 3 independent experiments as shown in panels F-G. **(I)** Fluorometric determination (arbitrary units) of ROS in the indicated cell lines as in F. Bar graphs are mean ± SEM, \**P*<0.05, \*\**P*<0.01, \*\*\**P*<0.001; n.s., not significant.

### The YAP/TAZ/TEAD transcription factors support PC expression in EmC

Analysis of *PC* mRNA levels in cells cultured in suspension under identical conditions as EmC but without rocking revealed lower levels similar to cells cultured in 2D, which suggested that mechanosignaling may be responsible for PC upregulation (**Supplemental Figure S3A**). The YAP and TAZ transcription factors are well-known mediators of mechano-signaling that are sequestered in the cytoplasm when inactive and do not bind DNA directly but rather through interaction with TEAD factors which are regulated through co-factor interactions (37, 38). E-cadherin-depleted cells showed reduced levels of total YAP protein and increased inactivating phosphorylation of YAP on Serine 127 (**Figure 4A**). To further explore the role of these transcription factors, we tested the effect of IK-930 which disrupts the interaction of YAP/TAZ with TEAD and therefore acts as a broad inhibitor of these transcription factor complexes (39). Treatment of established emboli with IK-930 downregulated PC protein and mRNA and expression (**Figures 4B-C**). The established YAP/TAZ target gene *CCN2* encoding CTGF (40) was used as a positive control. Transient silencing of *YAP* was sufficient to also downregulate *PC* and *CCN2* (**Figure 4D**). PC downregulation and silencing of YAP were confirmed at the level of protein, which also showed that depletion of YAP did not affect the level of E-cadherin (**Figure 4E**). Consistent with the reduction of PC expression, mitochondrial respiration was compromised upon IK-930 treatment (**Figures 4F-G**) or knockdown of YAP expression (**Figures 4 H-I**). Either treatment reduced the number of live cells recovered from EmC, though not to the same extent as loss of E-cadherin (**Supplemental Figures S3B-E**).

**Figure 4.**
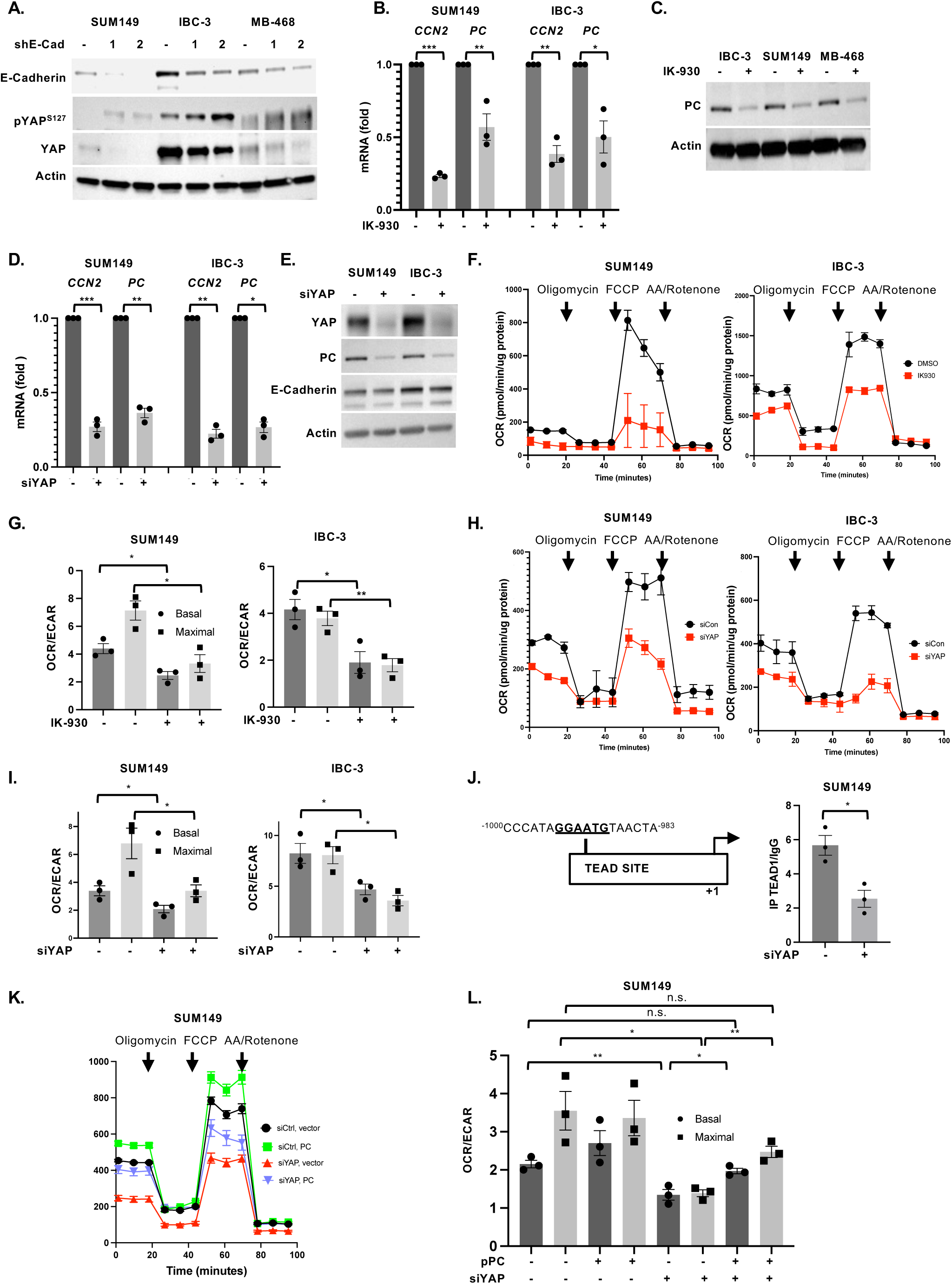
The YAP/TAZ/TEAD transcription factors support PC expression in EmC. **(A)** Western analysis of YAP and pYAP^S127^ levels in the indicated cells lines with stable E-cadherin depletion and controls. **(B)** qRT-PCR of *CCN2* and *PC* expression in SUM149 and IBC-3 treated with IK-930 (5 µM) for 3 days, similar to schematic in Figure 3A. Normalized values are shown as fold-change compared to controls. **(C)** Western analysis of PC expression in cells as in panel B. Actin served as a loading control. **(D)** qRT-PCR of *CCN2* and *PC* expression in SUM149 and IBC-3 after transient transfection with siRNA against *YAP*. Normalized values are shown as fold-change compared to controls. **(E)** Western analysis of cells as in panel D. **(F)** Oxygen consumption rate (OCR) over time of SUM149 and IBC-3 cells treated with -/+ IK-930 (5 µM) for 12 h. **(G)** OCR/ECAR ratios of cells as in panel F (n=3). **(H)** Oxygen consumption rate (OCR) over time of SUM149 and IBC-3 after transient transfection with siRNA against *YAP*. **(I)** OCR/ECAR ratios of cells as in panel H (n=3). **(J)** Schematic representation of the TEAD binding site in the PC promoter and ChIP analysis for TEAD to this element in cells with or without YAP depletion by siRNA. **(K)** Representative Seahorse analysis of the oxygen consumption rate (OCR) over time from SUM149 cells with siYAP and PC overexpression as indicated. **(L)** Basal and maximal OCR/ECAR ratios of cells from 3 independent experiments as shown in panel K. Bar graphs show mean ± SEM, \**P*<0.05, \*\**P*<0.01, \*\*\**P*<0.001; n.s., not significant.

Analysis of the *PC* promoter indicated a consensus TEAD binding site sequence about 1 kb upstream of the transcription start site, and chromatin immunoprecipitation indicated that TEAD binding was compromised when YAP expression was silenced in SUM149 cells (**Figure 4J**). Accordingly, the activity of a *PC* promoter-reporter construct was significantly reduced in YAP-depleted cells (**Supplemental Figure S3F**). To test whether PC functions downstream of YAP, we overexpressed PC in YAP-depleted cells which was sufficient to restore basal OCR and OCR/ECAR ratios (**Figures 4K-L**). Collectively these data show that *PC* is a target gene of YAP/TEAD and that YAP operates downstream of E-cadherin to promote mitochondrial respiration through PC.

### E-cadherin activates YAP through the AKT pathway

E-Cadherin can interact with receptor tyrosine kinases such as EGFR and either inhibit or activate downstream signaling pathways such as PI3K/AKT (12), which were similarly reported to activate or inhibit the YAP/TAZ transcription factors (41, 42). Analysis of E-Cadherin-depleted cells showed reduced AKT autophosphorylation as a measure of its activation (**Figure 5A**). When established emboli were treated with the AKT inhibitor MK2206 or PI3K inhibitor LY2940002, *PC* mRNA expression was reduced along with the YAP-target *CCN2* (**Figure 5B**). In addition, downregulation of PC and YAP by AKT inhibition was validated at the level of protein (**Figure 5C**). These data are consistent with the reported ability of AKT to prevent YAP degradation (43). Furthermore, AKT inhibition decreased mitochondrial respiration (**Figures 5D-E**). Consistent with AKT being upstream of several targets that may promote survival and/or proliferation, treatment of established emboli with MK2206 also reduced the number of viable cells (**Supplemental Figures S4A-B**). In summary, these data suggest that E-Cadherin regulates PC through stabilization and activation of YAP signaling.

**Figure 5.**
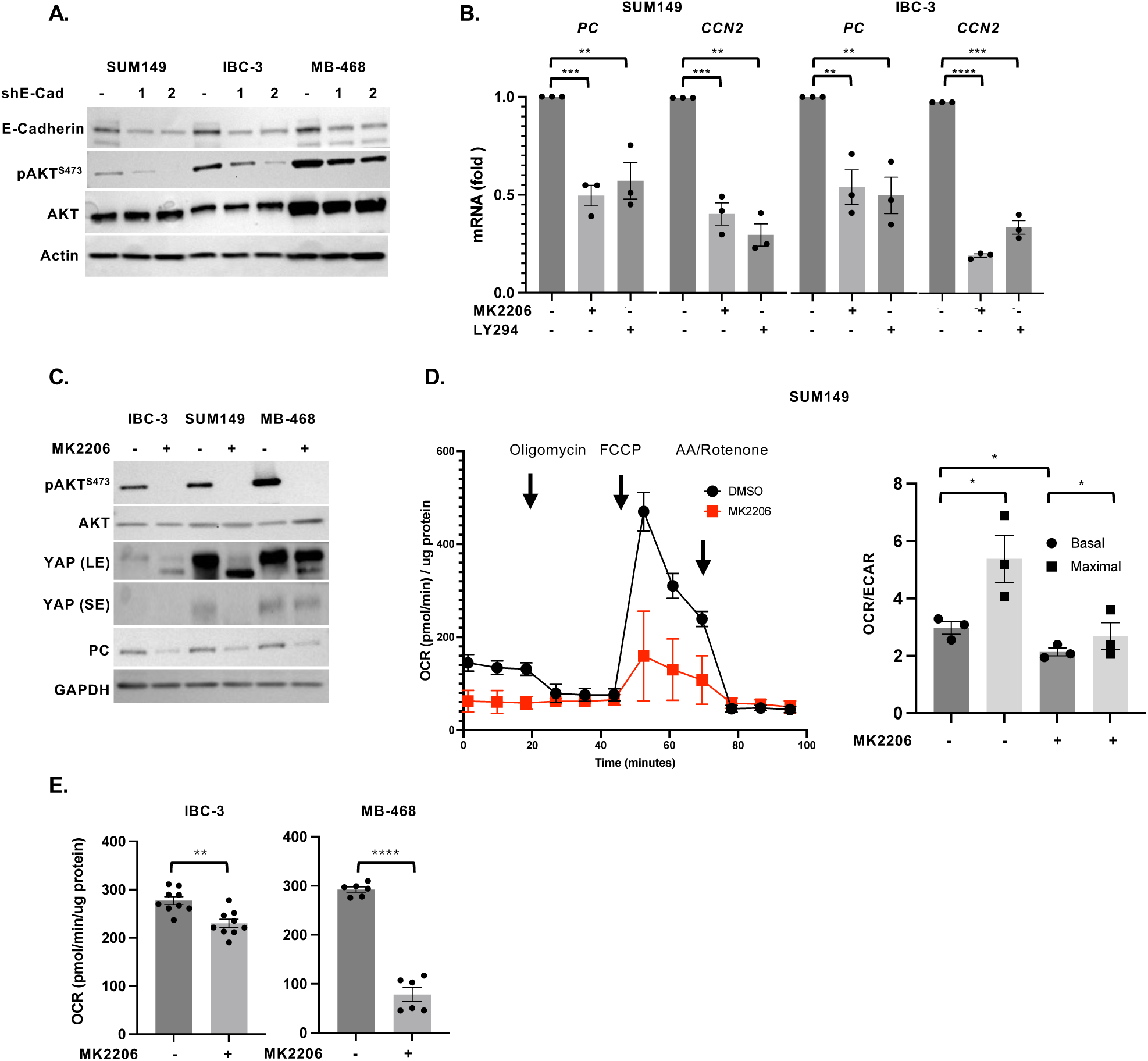
E-Cadherin activates YAP through AKT. **(A)** Western analysis of indicated proteins in SUM149, IBC-3, and MB-468 cells with stable depletion of E-cadherin and control. **(B)** qRT-PCR analysis of *PC* and *CCN2* mRNA levels in SUM149 and IBC-3 cells treated for 3 days with the AKT inhibitor MK2206 or PI3K inhibitor LY294002 (similar to schematic in Figure 3A). **(C)** Western analysis of cells as in panel B. **(D)** Oxygen consumption rate (OCR) over time in SUM149 cells treated with MK2206 for 12 h (similar to schematic in Figure 3B), and OCR/ECAR ratios from three independent experiments. **(E)** Basal OCR of IBC-3 and MB-468 cells as in panel D. Bar graphs show mean ± SEM, \**P*<0.05, \*\**P*<0.01, \*\*\**P*<0.001, \*\*\*\**P*<0.0001.

### ZY-444 attenuates growth of established primary tumors and experimental metastases

To test the effect of PC inhibition as monotherapy *in vivo*, we treated mice with well-established SUM149 tumors with ZY-444, which significantly attenuated tumor growth compared to DMSO treatment as control (**Figure 6A**, individual tumor growth curves and mouse body weights are shown in **Supplemental Figures S5A-B).** Distant metastases are the primary cause of death for BC patients, and experimental models suggest that lung and bone metastases specifically depend on OXPHOS (8). Therefore, we also assessed the effect of PC inhibition on the growth of established experimental lung metastases. Mice were injected via the tail vein with luciferase-expressing SUM149 cells and monitored for formation of lung colonies through bioluminescence imaging (BLI). When all mice showed signals above background, they were treated with ZY-444 or DMSO as control. Comparison of the bioluminescent signals at the beginning and end of treatment showed that in the control groups BLI signals increased significantly over time (**Figures 6B-C**). In the treatment group most mice showed attenuation or even regression of luminescent signal. At the endpoint, lungs were fixed and analyzed for lesions by image analysis (**Figure 6D**). Enumeration of lesions above 15,000 µm^2^ (**Figure 6E**) confirmed that the ZY-444 group exhibited significantly less tumor load (**Figure 6F)**. Collectively, these data show that monotherapy of mice with ZY-444 significantly attenuated the growth of an aggressive human tumor cell line within the orthotopic and lung environments. While higher doses or longer treatments could be assessed for grater efficacy, we also explored the potential for combination therapies. Because glutamine is another key metabolite that fuels the TCA cycle, we evaluated the effect of combining the glutamine antagonist DON (44) with ZY-444 on established EmC cultures. Across all the three cell lines the combination treatment reduced the number of viable cells retrieved from cell clusters compared to treatment with either drug alone (**Figures 6G** and **Supplemental Figure S5C**). MB-468 cells were most sensitive and IBC-3 least, suggesting that these cell lines are valuable models for investigating mechanisms of therapeutic response and metabolic resistance. Taken together, these findings suggest that dual inhibition of glutamine metabolism and PC may enhance anti-tumor efficacy as outlined in the proposed signaling model **(Figure 7).**

**Figure 6.**
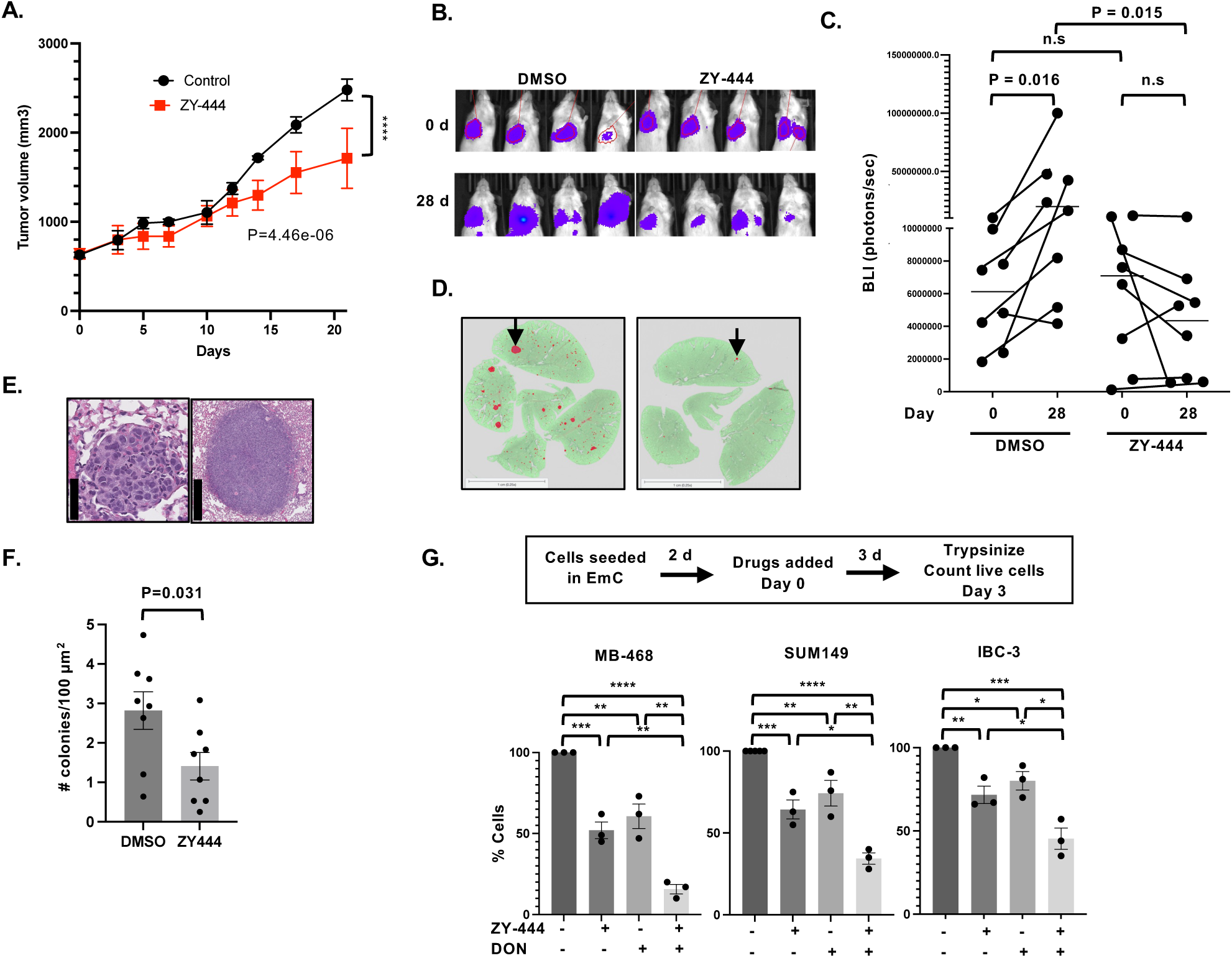
PC inhibition by ZY-444 attenuates growth of primary tumors and experimental metastases. **(A)** Tumor volumes of established SUM149 xenografts over 21 days from initiation of treatment (d 0) with ZY-444 or DMSO as control (n=6-7). A linear mixed-effects model with Animal as a random effect was used to determine Day × Treatment interaction on log-transformed tumor volumes. **(B)** Illustration of bioluminescence imaging of lungs in mice before (0 d) and after 28 days of treatment with ZY-444 or DMSO as control. **(C)** Quantification of bioluminescence as shown in panel A, n=8. P values within groups comparing 0 and 28 d were determined by paired Wilcoxon signed rank exact test. Within day comparison of groups was analyzed by unpaired Wilcoxon signed rank sum exact test (a.k.a. Mann-Whitney Test). **(D)** Examples of lung sections with many(left), respectively few (right) SUM149 colonies, one each indicated by an arrow. The HALO-annotation mask of the lung is shown in green and lesions in red (scale bar = 1 mm). **(E)** Examples of H&E-stained lung colonies just above 15,000 µm^2^ (left, scale bar: 50 µm) and a large lesion (right, scale bar 500 µm). **(F)** Number of colonies per 100 mm^2^ lung area in four step sections per lung (n = 8 mice). (**G**) Schematic of the experimental timeline of EmC and treatment with ZY-444 (2.5 µM) and/or DON (5 µM) and quantification of % viable cells retrieved from emboli and aggregates compared to DMSO controls (n=3). Bar graphs show mean ± SEM, \**P*<0.05, \*\**P*<0.01, \*\*\**P*<0.001, \*\*\*\**P*<0.0001.

**Figure 7.**
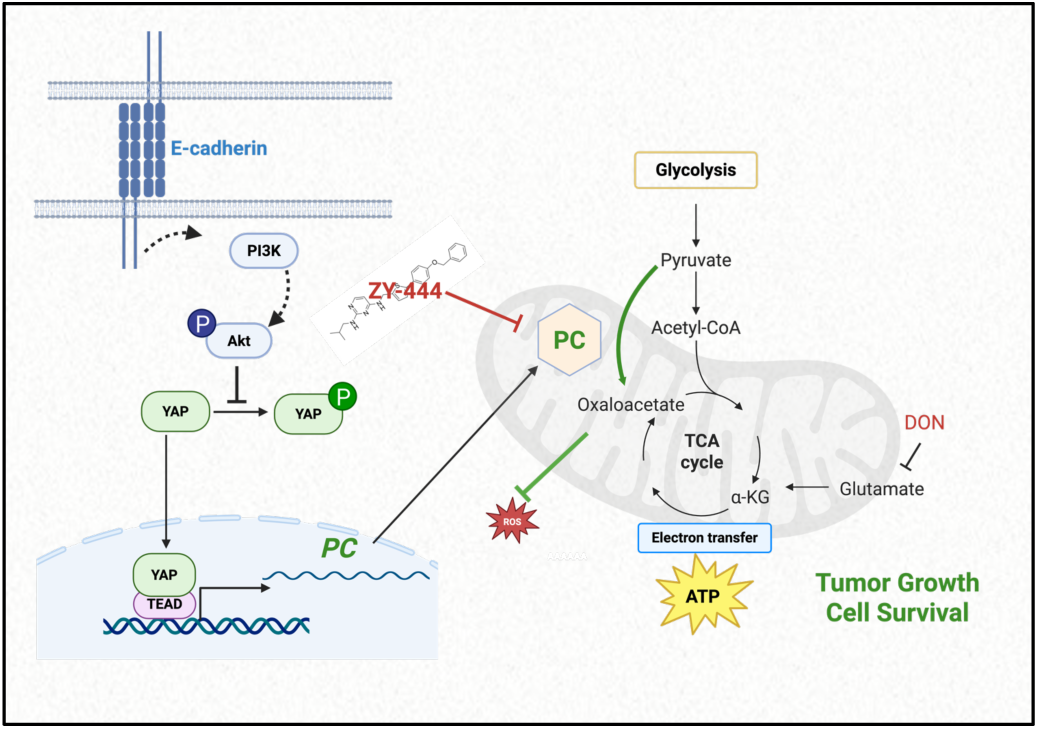
Schematic of the pathway elucidated in this report. Generated with BioRender.com.

## DISCUSSION

Crosstalk between cell-cell adhesion and metabolism has been recognized as a feature of cancer progression (45), but the mechanistic details and extent of this interaction remain to be fully elucidated. Our findings expand the understanding of E-cadherin’s pro-oncogenic role in cancer by identifying its contribution to mitochondrial metabolism through YAP-mediated expression of pyruvate carboxylase. Through this pathway, E-cadherin promotes oxidative phosphorylation while simultaneously reducing oxidative stress. This dual function explains in part why E-cadherin is retained or re-expressed in aggressive breast cancers, particularly in inflammatory breast cancer and certain triple-negative breast cancer subtypes. The requirement of E-cadherin for YAP-activation contrasts its well-characterized role as an inhibitor of YAP through its sequestration in the cytoplasm.

Many studies have investigated the potential of E-cadherin as a prognostic marker. Apart from loss of E-cadherin being diagnostic for the lobular breast carcinoma subtype, no association with outcome was found in a large study of multiple subtypes of ductal carcinomas (11). Others found that deviation of both lower and higher than median expression associated with poor prognosis (46) or that low expression associated with higher grade and worse outcomes in TNBC (47). In other words, E-cadherin is rarely completely lost even in aggressive breast cancers. Using isogenic pairs of E-cadherin high and low-expressing variants of the TNBC cell lines MDA-MB-468 and MDA-MB-231 in 2D cultures, Lee et al. showed that E-cadherin supports the serine synthesis pathway through expression of the rate liming enzyme PHGDH, diverting pyruvate away from the TCA cycle which is then supported by glutamine instead (18). While this pathway was not specifically induced by EmC (24), our approach showed that 3D culture in EmC further increased oxidative phosphorylation of these cells through E-cadherin-mediated upregulation of PC. PHGDH is most highly expressed in estrogen-receptor negative breast cancers (48). Our results suggest that in vivo PC may cooperate with the serine pathway to maximizing the utilization of pyruvate and further combat oxidative stress through the malate anti-oxidant system (36).

The identification of the E-cadherin→AKT→YAP/TEAD→PC signaling pathway revealed the convergence of adhesion signaling, mechanotransduction, and metabolic regulation. The involvement of YAP/TEAD transcription factors is particularly intriguing given their well-established role in other contexts in promoting EMT and mesenchymal phenotypes as well as glycolysis (49–53) and to inhibit oxidative mitochondrial metabolism (54). However, by generating pyruvate, glycolysis is intimately linked to OXPHOS, and these pathways are therefore anything but mutually exclusive (55). Our data provide context for the functional output of YAP/TEAD activity. In the presence of E-cadherin-mediated cell-cell adhesion and under mechanical stimulation, YAP/TEAD drives an epithelial metabolic program through PC expression and OXPHOS, rather than promoting purely mesenchymal traits. The direct binding of TEAD to the PC promoter, confirmed by our ChIP experiments, establishes PC as a *bona fide* YAP/TEAD target gene, expanding the known target gene repertoire of these factors.

The contribution of mechanical stimulation as experienced in EmC to PC induction suggests that YAP/TEAD integration of both biochemical (E-cadherin/AKT) and biophysical (mechanical) signals, both potentially mediated by E-cadherin, is necessary for full activation of the PC transcriptional program. Traditionally, E-cadherin had been considered to be a tumor suppressor because it attenuates cancer cell motility and mediates contact inhibition of cell proliferation in part through sequestration of YAP in the cytoplasm and through stimulation of its degradation (56, 57). One study reported that mechanical strain on E-cadherin adhesions in monolayer cultures of MDCK cells releases YAP from cytoplasmic sequestration followed by nuclear translocation. In this case, the extracellular domain of E-cadherin was required for cytoplasmic retention of YAP (58) suggesting the involvement of interaction with other membrane spanning proteins in the regulation of YAP activity. E-cadherin can associate with many transmembrane signaling molecules. For example, interaction with the EGF receptor system can lead to inhibition of ligand-dependent signaling but also to activation of ligand-independent signaling which is in part modulated by mechanical tension (59–62). Receptor tyrosine kinases activate the PI3K/AKT pathway, which in turn can activate the TAP/TAZ/TEAD transcription factors in part by preventing their degradation (43, 63). E-cadherin may contribute to AKT/YAP activation through transmembrane signaling molecules and also contribute to the mechanosensing through PI3K (42). Cell-cell adhesions experience tensions even in normal tissue due to the pulsatile flow of blood and lymph and normal body movements the contribution of which was proposed as a mechanism for lung cancer progression (64).

Targeting of OXPHOS in metastasis and therapy resistance is a relatively novel frontier in cancer therapy (19, 65–67). Our work has revealed an important molecular mechanism underlying the metabolic characteristic of epithelial BC. Pyruvate carboxylase is implicated in cancer progression by providing metabolic flexibility and protection from oxidative stress (29). Breast cancers, but not normal breast epithelium, express PC (31, 34). In mouse models, PC has been particularly associated with the promotion of BC lung metastasis (28–32). PC expression promotes colonization of the oxygen-rich lungs, while PC expression in primary tumors may support immune surveillance due to reduced lactate content (31, 68). In the 4T1 model, systemic pharmacological inhibition of PC with ZY-444 from the start of tumor onset inhibited metastasis but also primary tumor growth (32), which confounds conclusions on the effect of ZY-444 on metastases. Additionally, inhibition of experimental lung colonization by ZY-444 (32) similarly cannot address the role of PC in established metastases. Therefore, we tested and showed the efficacy of ZY-444 in attenuating the growth of both established orthotopic tumors and lung colonies, the clinically most relevant paradigm. ZY-444 is well tolerated in mice (32, 69), suggesting that PC inhibition could have potential utility in the clinic. The development of more potent and specific PC inhibitors could provide new therapeutic options for metastatic breast cancer patients whose tumors retain epithelial characteristics, and warrant studies of PC intersection with other metabolic and signaling pathways for combination therapies, such as by inhibition of glutamine availability (18, 70, 71).

The identification of the E-cadherin-AKT-YAP/TEAD-PC pathway offers multiple potential therapeutic intervention points. Our demonstration that clinically relevant AKT inhibitors (MK2206) and TEAD inhibitors (IK-930) reduce PC expression and mitochondrial respiration suggests that existing drugs targeting these pathways may exert part of their anti-cancer effects through metabolic reprogramming. This has important implications for patient stratification and combination therapy design. E-cadherin-positive breast cancers, including inflammatory breast cancer and basal-like triple-negative breast cancers, may be particularly sensitive to therapies targeting this pathway. Furthermore, the co-expression of E-cadherin and PC in patient specimens could serve as a biomarker to identify tumors likely to respond to PC inhibition or upstream pathway blockade. Additionally, given the role of this pathway in protecting against oxidative stress, combination strategies pairing PC inhibition with ROS-inducing agents may show synergistic efficacy.

### Limitations and future directions

While our study establishes a clear mechanistic link between E-cadherin and PC-mediated metabolism, several questions remain. First, although our emboli culture system recapitulates key aspects of tumor cell aggregates in vivo, further studies in vivo will be necessary to fully validate the therapeutic potential of PC inhibition by higher doses and by combination therapies using additional models in the immunocompetent mouse. Secondly, the mechanisms by which E-cadherin activates AKT in 3D culture warrant deeper investigation, including identification of the relevant receptor tyrosine kinases and co-receptors involved. In addition, the full spectrum of YAP/TAZ/TEAD factors capable of inducing PC expression and their intersection with other potentially metabolically regulated transcription factors in breast cancer requires further attention. Third, our results suggest that this pathway could be particularly relevant for circulating tumor cell clusters which will be addressed in future studies.

## MATERIALS AND METHODS

### Sex as biological variable

This study exclusively used female mice because the cell lines were derived from women’s breast cancers and the disease modeled is most prevalent in women.

### Antibodies and reagents

Antibodies were obtained from the following sources: Cell Signaling Technology (E-cadherin (24E10), #3195; pyruvate carboxylase, #66470; YAP/TAZ, #8418; YAP, #4912S; pYAP^S127^, #13008S; pAKT^S473^, #4060; AKT, #9272; Santa Cruz Biotechnology (GAPDH, #sc-47724); and DSHB (A-Actin, #JLA20). IK-930 (#E1483), MK2206 (#S1078), LY294002 (#S1105) were from Selleck chemicals. ZY-444 (#TA9H998A31C1), 6-Diazo-5-oxo-L-norleucine (DON) (#D2141) and DMSO (#D-2650) were from MilliporeSigma. DMSO was used as a vehicle volume control in all experiments involving drug treatments.

### Cell culture and drug treatments

SUM149 and SUM159 originated from Asterand Bioscience; MDA-MB-468 cells were from ATCC. IBC-3 and MDA-MB-231-LM2 were kind gifts from Drs. Wendy Woodward (MDACC) and J. Massagué (MSKCC), respectively. Cells were used at passages 2-30, authenticated last in 2025 by GenePrint®10 (Promega), and tested annually for *Mycoplasma* by qPCR. Cells were cultured at 5% CO_2_/37°C in media with 100 units/ml penicillin and 100 µg/ml streptomycin as follows: MDA-MB-468 and MDA-MB-231-LM2 in Dulbecco’s modified Eagle’s medium (DMEM, GIBCO #11965118); SUM149, and IBC-3 in Ham’s F-12 media (GIBCO, #31765092) with 5 μg/ml hydrocortisone (MilliporeSigma #H-0135) and 1 μg/ml Insulin (MilliporeSigma #I-0516); SUM159 in RPMI (Quality Biologicals #112-024-101) with 2 mM glutamine (GIBCO #25030081), 10 mM HEPES (GIBCO #15630080), 1 mM sodium pyruvate (GIBCO #11360070), 1X nonessential amino acids (GIBCO, #11140-050) and 55 μM β-mercaptoethanol (GIBCO, #21985-023); In addition, all media contained 2.25% PEG8000 (MilliporeSigma #202452) as per emboli culture (EmC) protocol (24). Unless specified otherwise, all data were derived from cells in EmC. Briefly, 1-2.5×10^5^ cells were seeded in 6-well ultra-low attachment (ULA) plates (Corning, #3471) in medium containing 2.25% PEG8000 on an orbital shaker at approximately 40 rpm for 2-5 days as indicated. Unless indicated otherwise, all analyses of 3D cultures were conducted after 3 days. Emboli were separated from single cells and small clusters by centrifugation at 500 rpm for 30 sec and washed once with PBS. For dissociations, emboli were treated for 10 min with TrypLExpress (GIBCO, #12604-013) which was then neutralized with cell culture medium. Cells were counted with a Countess (Life Technologies, USA) using trypan blue dye exclusion. For assessment of drug effects, drugs were added after 2 days of EmC, and emboli collected 3 days later.

### Generation of cells with lentiviral shRNA-mediated E-Cadherin knockdown

The shRNAs against *CDH1* encoding E-cadherin were from Millipore SIGMA (TRCN0000237841, shCDH1#1, 5’-AGATTGCACCGGTCGACAAAG-3’; TRCN0000237842, shCDH1#2, 5’-TTTCGGCAGTTCAAGCTATAT-3’). The shRNAs were inserted into the pLKO.1 vector (SIGMA, #SHC016) and transfected into HEK-293T cells along with lentiviral packaging mix (SIGMA, #SHP001) using PolyJet in vitro DNA transfection reagent (SignaGen Laboratories, #SL100688). After 48 h, medium from the transfected cells was collected, centrifuged, and filtered. 0.5 ml viral supernatant media, supplemented with 10 μg/ml polybrene, were added to the target cells. Pools of stable transfectants were selected by culture with 1 μg/ml puromycin for at least 1 week.

### Western blotting analysis

Western blotting was carried out according to standard procedures. Briefly, cells were lysed for 15 minutes on ice with cell lysis buffer containing 10 mM Tris, 1 mM EDTA, 400 mM NaCl, 0.1% NP-40, 10 μL/mL each of protease inhibitor cocktail (MilliporeSigma #P8340), phosphatase inhibitor cocktail #2 (MilliporeSigma, P5726), and phosphatase inhibitor cocktail #3 (MilliporeSigma, P0044). Samples were centrifuged at 12,700 rpm for 15 min at 4°C. The amounts of total protein were quantified by BCA assay (Thermo Fisher Scientific, 23225) for equal loading. The loaded samples were electrophoresed by SDS-PAGE and transferred to Nitrocellulose membranes. After blocking with 5% non-fat dry milk solution for 1 h, the membranes were incubated with primary antibodies overnight, washed 3X10 min each with TBST (TBS containing 0.05% Tween-20). Appropriate secondary antibodies were added for 1 h followed by washing the membrane before scanned by an iBRIGHT 1500 Imaging System (Thermo Fisher Scientific).

### RNA isolation and quantitative RT-PCR

RNA was isolated using GeneJET RNA purification kit (Thermo Scientific, #K0732) and 2 µg RNA was taken for cDNA synthesis using Superscript Reverse Transcriptase III (RT) according to the manufacturer’s instructions (Invitrogen, #18080044). PCR was carried out with Fast SYBR Green master mix (#4385612, Applied Biosystems, Foster City, USA) and the Quantstudio 5 Real**-**Time PCR instrument (Applied Biosystems). Relative expression levels were measured using the relative quantitation ΔΔ*C*t method normalized to RPLPO. Data are from three independent biological replicates, each assayed in triplicates. Primers were as follows, *<u>PC</u>* <u>(5’-ccagaggcaggtcttctttg-3’ and 5’-gggtgaggtcaccacagtct-3’); *CCN2* (5’-taccaatgacaacgcctcct-3’ and 5’-ccgtcggtacatactccaca-3’);</u> *RPLP0* (5’-gcaatgttgccagtgtctgtc-3’ and 5’-gccttgaccttttcagcaagt-3’).

### Transient transfections and luciferase reporter assays

For silencing of *YAP1* by a pool of siRNA (sc-38637) and/or overexpression of pyruvate carboxylase (Addgene #184466) or E-Cadherin (72), a kind gift from Dr. Deborah Morrison, NCI. 2 × 10^6^ cells were nucleofected using AMAXA nucleofection (Lonza, MD, USA), trypsinized 24 h later, and seeded onto 6 well ULA plates for EmC as indicated. Silencing and overexpression were confirmed as shown in respective figures. For promoter-reporter assays, 2 million cells were nucleofected with 20 nM siRNA oligos along with 5 μg luciferase reporter plasmid containing 1108 bp of the *PC* promoter P2 (73), a kind gift from Dr. Jitrapakdee, Thailand, or vector control. After 24 h, transfected cells were trypsinized, counted, and 250,000 cells were seeded onto 6 well ULA plates for EmC culture. Three days later, cell extracts were prepared and luciferase activity assayed using the Luciferase Reporter Assay kit (Promega, #E1910).

### Chromatin immunoprecipitation (ChIP) assay

SUM149 cells were transfected with control or *YAP1* siRNA and 24 h later, cells were trypsinized, counted and cultured in EmC for 3 days. ChIP analysis was performed as per the manufacturer’s instructions (EZ ChIP, #17-371, MilliporeSigma, USA). Briefly, cells were cross-linked and chromatin was prepared and sonicated to an average size of 500 bp. The DNA fragments were immunoprecipitated with 5 µg antibodies specific to TEAD1 (Cell Signaling Technology, #12292T) or control IgG (Thermo Fisher Scientific, #31235) at 4°C overnight. After reversal of the cross-linking, the immunoprecipitated chromatin was taken for qRT-PCR of the *PC* promoter region with forward (5’-ACTACCTACTCAGAGACATGTCA-3’) and reverse (5’-ATG AGG GGA AGG CAT GTA GG-3’).

### Seahorse Cell Mito Stress test

The oxygen consumption rate (OCR) and extracellular acidification rate (ECAR) were measured using a Seahorse XF-96 analyzer with the Seahorse XF Cell Mito Stress Test kit (Cat#103015-100, Seahorse Bioscience, MA, USA) according to the manufacturer’s instructions. Cells from EmC were trypsinized for 10 min using TRYPLExpress, counted, washed with Seahorse Bioscience XF assay medium (pH 7.4) supplemented with 1 mM pyruvate, 2 mM glutamine, and 10 mM glucose and 200,000 cells were seeded onto Seahorse 96-well microplates coated with Poly-L-Lysine. The plate was placed in a 37 °C incubator without CO_2_ for 1 h before transfer to the Seahorse XF-96 analyzer. Oligomycin (1 μM), protonophoric uncoupler (FCCP, 2 μM), and antimycin A + rotenone (0.5 μM each) were preloaded in reagent Ports A, B, and C. OCR and ECAR values were automatically calculated and recorded by the sensor cartridge and Seahorse XF-96 software and were normalized to the protein amount in each well. For calculation of OCR/ECAR ratios, the mean OCR from each experiment was divided by the respective mean ECAR. Three-day cultures were used unless indicated otherwise. As applicable, drug treatments were started after 2.5 days for a duration of 12 h.

### Measurement of oxidative stress

The Oxidative Stress Indicator CM-H2DCFDA (Thermo Fisher Scientific, #C6827) was used to evaluate the level of reactive oxygen species (ROS). Briefly, 50,000 cells were resuspended in 400 µl serum-free medium, followed by addition of 5 µM CM-H2DCFDA for 20 min in the dark at 37°C. After 20 min, 400 µl ice-cold PBS + 0.5% BSA were added to stop the reaction, the cells were pelleted, washed twice, resuspended in 200 µl PBS + 0.5% BSA, and transferred to a 96-well plate (Greiner, #655090). The DCF fluorescence was measured with a Spectramax iD3 plate reader (Molecular Devices, USA) using excitation (492 nm) and emission wavelengths (535 nm) settings.

### Animal studies and experimental metastasis assay

Animal care was provided in accordance with the procedures outlined in the Guide for the Care and Use of Laboratory Animals (National Academies Press, 2011) including those pertaining to studies of neoplasia (National Research Council, 1996). All experiments were conducted under protocols approved by the IACUC at NCI-Frederick. NCI-Frederick is accredited by Association for Assessment and Accreditation of Laboratory Animal Care International (AALACi) and follows the Public Health Service Policy for the Care and Use of Laboratory. For experimental metastasis assays, cells were dissociated from EmC and 1×10^6^ cells in 100 μl PBS were injected intravenously by tail vein into 8-12-week-old female NSG mice. Within 2 h of injections, bioluminescence was measured using the Xenogen IVIS to verify inoculation of the lungs. Bioluminescence was monitored at two-week intervals and treatments began when BLI was at least 100,000 photons/sec above background. Mice were injected i.p. with either vehicle (6.85% DMSO in water) or ZY-444 (5 mg/kg) on Mondays, Wednesdays, and Fridays for 4 weeks (day 28). Upon termination, the lungs were inflated with 10% neutral-buffered formalin, and lobes were separated before embedding. Four 5 μm step sections (100 μm apart, 2 per slide), were stained with H&E. For the histologic confirmation by a board-certified veterinary pathologist and quantification of lung metastases, slides were scanned with an Aperio AT2 scanner (Leica Biosystems) at 20x objective (0.50um/pixel) and analyzed with HALO software (Indica Labs) to quantify metastases and total lung area for normalization. Lesions with >15,000 μm^2^ tumor cell area were validated by a veterinarian pathologist blinded to the experiment. For the enumeration of colonies, care was taken not to double-count lesions that were represented on more than on section. For primary tumor studies, 2×10^6^ SUM149-GFP-Luc cells in 50 μl DMEM were surgically implanted into the inguinal mammary fat pad of 20-22-week-old female NSG mice. When tumors reached 500-800 mm^3^ tumor volume (day 0) mice were treated with ZY-444 or DMSO as above for 3 weeks (day 21). CTCs were analyzed by IMC as described (74).

### Tissue microarray (TMA), multiplex immunofluorescence and image analysis

Human FFPE breast cancer tissue microarray BR1505e was purchased from TissueArray.Com LLC. Staining of 5 μm section was performed using a Leica Bond RX autostainer (Leica Biosystems). Initial antigen retrieval was performed using EDTA buffer for 20 min at 100℃ on the Bond autostainer. OPAL^TM^ multiplex staining was accomplished with the following antibodies dilutions, incubation times and reporter dyes, applied in this order: (1) E-Cadherin (Cell Signaling #3195), 1:200, 30 min, Opal690 (Akoya Biosciences FP1497001KT); (2) Pyruvate carboxylase (Protein Tech Group #16588-1-AP), 1:1000, 60 min, Opal570 (Akoya Biosciences # FP1488001KT); (3) Vimentin (Abcam #ab92547), 1:2000, 30 min, Opal520 (Akoya Biosciences #FP1487001KT). Primary antibody detection was accomplished using the Bond Polymer Refine Detection kit (Leica Biosystems #DS9800) with the Post-Primary reagent, DAB, and Hematoxylin removed from Leica’s default staining protocol. Reporter dyes were added to the tissue per the manufacturer’s instructions. Between stainings, EDTA antigen retrieval solution was applied again to the tissues for 20 min at 95 ℃. All slides were counterstained with DAPI and then digitally scanned at 20x using an Akoya PhenoImager HT whole-slide scanner (Akoya Biosciences), followed by spectral unmixing to resolve fluorophore signals. Individual tissue cores were evaluated for suitability for analysis and manually annotated to exclude artifacts. Cell-based quantification of immunofluorescence was performed using the Cytonuclear FL algorithm (v2.0.12) within HALO AI (v4.2; Indica Labs) to classify cellular positivity for each marker. Cellular phenotypes were defined based on expression of vimentin, pyruvate carboxylase, and E-cadherin. Data are from 18 hormone receptor negative specimen (9 TNBC, 9 HER2+) and 38 estrogen receptor positive specimen, one core each chosen from duplicates by based on total number of cells (>1000) expressing at least one of three markers.

### Statistics

Unless stated otherwise, quantitative data are shown as mean ± S.E.M. and were analyzed by the two-tailed unequal variance t-test. The number of samples (n) refers to biological replicates. For the tumor growth time-series, data were analyzed using a linear mixed-effects model implemented in the *lme4* R package (v1.1.37) (75). Animal was included as a random effectand tumor volume was log-transformed prior to analysis. P-values were estimated using the *lmerTest* R package (v3.1.3) (76). Fixed effects included Treatment, Day, and their interaction.

## Data availability

Data are available in public repositories (no large datasets were generated), in the Supporting Data Values file, or from the corresponding authors upon request.

## AUTHOR CONTRIBUTIONS

The contributions of the authors were as follows: conceptualization (KB, ES), investigation (KB, JMW, LM, BAG, DOB), resources (KB, LM, SS, SJL, DWM, ES), data curation (KB, JMW, DAS, AFE, DD, LB, ES), formal analysis (KB, JMW, DAS, AFE, DD, LB, ES), validation (KB, JMW, LM, AFE, DD, LB, ES), visualization (KB, JMW, LB, ES), methodology (KB, JMW, SS, BAG, DOB, DAS, AFE, ES), writing of original draft (KB, ES), reviewing/editing (KB, SS, BAG, DOB, DAS, AFE,LB, DWM, ES), project administration (KB, ES), supervision (KB, SJL, DWM, ES), funding acquisition (SJL, DMW, ES).

## ACKNOWLEDGEMENTS

We are grateful for the superb support provided by the Laboratory Animal Sciences Program including Small Animal Imaging Program and the Molecular Histotechnology Laboratory of the Frederick National Laboratory for Cancer Research, and to Dr. Milind Pore and Ashley Cardamone for IMC analysis of CTCs. We also thank MaryBeth Hilton and Dr. Brad St. Croix (NCI/CCR) for their kind assistance with NSG mice.

This research was supported by the Intramural Research Program of the National Institutes of Health (NIH), ZIA BC 010307, and in part with federal funds under contract no. 75N91019D00024, and is subject to the NIH Public Access Policy. Through acceptance of this federal funding, the NIH has been given a right to make the work publicly available in PubMed Central. The contributions of the NIH author(s) were made as part of their official duties as NIH federal employees, are in compliance with agency policy requirements, and are considered Works of the United States Government. However, the findings and conclusions presented in this paper are those of the author(s) and do not necessarily reflect the views of the NIH or the U.S. Department of Health and Human Services, nor does mention of trade names, commercial products, or organizations imply endorsement by the U.S. Government.

## REFERENCES

1. Yang J, Antin P, Berx G, Blanpain C, Brabletz T, Bronner M, et al. Guidelines and definitions for research on epithelial-mesenchymal transition. Nat Rev Mol Cell Biol. 2020;21(6):341–52.

2. Brabletz S, Schuhwerk H, Brabletz T, and Stemmler MP. Dynamic EMT: a multi-tool for tumor progression. EMBO J. 2021;40(18):e108647.

3. Jiang B, Mu Q, Qiu F, Li X, Xu W, Yu J, et al. Machine learning of genomic features in organotropic metastases stratifies progression risk of primary tumors. Nat Commun. 2021;12(1):6692.

4. Williams ED, Gao D, Redfern A, and Thompson EW. Controversies around epithelial-mesenchymal plasticity in cancer metastasis. Nat Rev Cancer. 2019;19(12):716–32.

5. Bruner HC, and Derksen PWB. Loss of E-Cadherin-Dependent Cell-Cell Adhesion and the Development and Progression of Cancer. Cold Spring Harb Perspect Biol. 2018;10(3).

6. Balamurugan K, Poria DK, Sehareen SW, Krishnamurthy S, Tang W, McKennett L, et al. Stabilization of E-cadherin adhesions by COX-2/GSK3beta signaling is a targetable pathway in metastatic breast cancer. JCI Insight. 2023;8(6):e156057.

7. Bornes L, Belthier G, and van Rheenen J. Epithelial-to-Mesenchymal Transition in the Light of Plasticity and Hybrid E/M States. J Clin Med. 2021;10(11).

8. Zuo Q, and Kang Y. Metabolic Reprogramming and Adaption in Breast Cancer Progression and Metastasis. Adv Exp Med Biol. 2025;1464:347–70.

9. Kowalski PJ, Rubin MA, and Kleer CG. E-cadherin expression in primary carcinomas of the breast and its distant metastases. Breast Cancer Res. 2003;5(6):R217–22.

10. Lim B, Woodward WA, Wang X, Reuben JM, and Ueno NT. Inflammatory breast cancer biology: the tumour microenvironment is key. Nat Rev Cancer. 2018;18(8):485–99.

11. Horne HN, Oh H, Sherman ME, Palakal M, Hewitt SM, Schmidt MK, et al. E-cadherin breast tumor expression, risk factors and survival: Pooled analysis of 5,933 cases from 12 studies in the Breast Cancer Association Consortium. Sci Rep. 2018;8(1):6574.

12. Rodriguez FJ, Lewis-Tuffin LJ, and Anastasiadis PZ. E-cadherin’s dark side: possible role in tumor progression. Biochim Biophys Acta. 2012;1826(1):23–31.

13. Hugo HJ, Gunasinghe N, Hollier BG, Tanaka T, Blick T, Toh A, et al. Epithelial requirement for in vitro proliferation and xenograft growth and metastasis of MDA-MB-468 human breast cancer cells: oncogenic rather than tumor-suppressive role of E-cadherin. Breast Cancer Res. 2017;19(1):86.

14. Kleer CG, van Golen KL, Braun T, and Merajver SD. Persistent E-cadherin expression in inflammatory breast cancer. Mod Pathol. 2001;14(5):458–64.

15. Chu K, Boley KM, Moraes R, Barsky SH, and Robertson FM. The paradox of E-cadherin: role in response to hypoxia in the tumor microenvironment and regulation of energy metabolism. Oncotarget. 2013;4(3):446–62.

16. Padmanaban V, Krol I, Suhail Y, Szczerba BM, Aceto N, Bader JS, et al. E-cadherin is required for metastasis in multiple models of breast cancer. Nature. 2019;573(7774):439–44.

17. Hapach LA, Carey SP, Schwager SC, Taufalele PV, Wang W, Mosier JA, et al. Phenotypic Heterogeneity and Metastasis of Breast Cancer Cells. Cancer Res. 2021;81(13):3649–63.

18. Lee G, Wong C, Cho A, West JJ, Crawford AJ, Russo GC, et al. E-cadherin Induces Serine Synthesis to Support Progression and Metastasis of Breast Cancer. Cancer Res. 2024.

19. Pendleton KE, Wang K, and Echeverria GV. Rewiring of mitochondrial metabolism in therapy-resistant cancers: permanent and plastic adaptations. Front Cell Dev Biol. 2023;11:1254313.

20. Li Z, Munim MB, Sharygin DA, Bevis BJ, and Vander Heiden MG. Understanding the Warburg Effect in Cancer. Cold Spring Harb Perspect Med. 2025;15(12).

21. Uslu C, Kapan E, and Lyakhovich A. Cancer resistance and metastasis are maintained through oxidative phosphorylation. Cancer Lett. 2024;587:216705.

22. Passaniti A, Kim MS, Polster BM, and Shapiro P. Targeting mitochondrial metabolism for metastatic cancer therapy. Mol Carcinog. 2022;61(9):827–38.

23. Whitaker-Menezes D, Martinez-Outschoorn UE, Flomenberg N, Birbe RC, Witkiewicz AK, Howell A, et al. Hyperactivation of oxidative mitochondrial metabolism in epithelial cancer cells in situ: visualizing the therapeutic effects of metformin in tumor tissue. Cell Cycle. 2011;10(23):4047–64.

24. Balamurugan K, Mikolaj MR, Weiss JM, Holewinski RJ, Fan Y, Dash SP, et al. 3D "emboli" culture models epithelial breast cancer cell oxidative mitochondrial metabolism with relevance for lung metastasis. Cancer Res Commun. 2026; :2025.08.24.671623.

25. Nolfi-Donegan D, Braganza A, and Shiva S. Mitochondrial electron transport chain: Oxidative phosphorylation, oxidant production, and methods of measurement. Redox Biol. 2020;37:101674.

26. Fogg VC, Lanning NJ, and Mackeigan JP. Mitochondria in cancer: at the crossroads of life and death. Chin J Cancer. 2011;30(8):526–39.

27. Chen K, Wang B, Shu H, Lyu J, Cui W, and Fang H. Oxidative phosphorylation at the crossroads of cancer: Metabolic orchestration, stromal collusion, and emerging therapeutic horizons. Interdisciplinary Medicine. 2025;3(6).

28. Hughey CC, and Crawford PA. Pyruvate Carboxylase Wields a Double-Edged Metabolic Sword. Cell Metab. 2019;29(6):1236–8.

29. Kiesel VA, Sheeley MP, Coleman MF, Cotul EK, Donkin SS, Hursting SD, et al. Pyruvate carboxylase and cancer progression. Cancer Metab. 2021;9(1):20.

30. Christen S, Lorendeau D, Schmieder R, Broekaert D, Metzger K, Veys K, et al. Breast Cancer-Derived Lung Metastases Show Increased Pyruvate Carboxylase-Dependent Anaplerosis. Cell Rep. 2016;17(3):837–48.

31. Shinde A, Wilmanski T, Chen H, Teegarden D, and Wendt MK. Pyruvate carboxylase supports the pulmonary tropism of metastatic breast cancer. Breast Cancer Res. 2018;20(1):76.

32. Lin Q, He Y, Wang X, Zhang Y, Hu M, Guo W, et al. Targeting Pyruvate Carboxylase by a Small Molecule Suppresses Breast Cancer Progression. Adv Sci (Weinh). 2020;7(9):1903483.

33. Sterneck E, Poria DK, and Balamurugan K. Slug and E-Cadherin: Stealth Accomplices? Front Mol Biosci. 2020;7:138.

34. Phannasil P, Thuwajit C, Warnnissorn M, Wallace JC, MacDonald MJ, and Jitrapakdee S. Pyruvate Carboxylase Is Up-Regulated in Breast Cancer and Essential to Support Growth and Invasion of MDA-MB-231 Cells. PLoS One. 2015;10(6):e0129848.

35. Rolver MG, Elingaard-Larsen LO, and Pedersen SF. Assessing Cell Viability and Death in 3D Spheroid Cultures of Cancer Cells. J Vis Exp. 2019(148).

36. Cappel DA, Deja S, Duarte JAG, Kucejova B, Inigo M, Fletcher JA, et al. Pyruvate-Carboxylase-Mediated Anaplerosis Promotes Antioxidant Capacity by Sustaining TCA Cycle and Redox Metabolism in Liver. Cell Metab. 2019;29(6):1291–305 e8.

37. Lin KC, Park HW, and Guan KL. Regulation of the Hippo Pathway Transcription Factor TEAD. Trends Biochem Sci. 2017;42(11):862–72.

38. Piccolo S, Panciera T, Contessotto P, and Cordenonsi M. YAP/TAZ as master regulators in cancer: modulation, function and therapeutic approaches. Nat Cancer. 2023;4(1):9–26.

39. Harvey KF, and Tang TT. Targeting the Hippo pathway in cancer. Nat Rev Drug Discov. 2025;24(11):852–69.

40. Lai D, Ho KC, Hao Y, and Yang X. Taxol resistance in breast cancer cells is mediated by the hippo pathway component TAZ and its downstream transcriptional targets Cyr61 and CTGF. Cancer Res. 2011;71(7):2728–38.

41. Basu S, Totty NF, Irwin MS, Sudol M, and Downward J. Akt phosphorylates the Yes-associated protein, YAP, to induce interaction with 14-3-3 and attenuation of p73-mediated apoptosis. Mol Cell. 2003;11(1):11–23.

42. Di-Luoffo M, Ben-Meriem Z, Lefebvre P, Delarue M, and Guillermet-Guibert J. PI3K functions as a hub in mechanotransduction. Trends Biochem Sci. 2021;46(11):878–88.

43. Jin Y, Xue Y, Yao J, Xu C, and Yu R. FBXO9 mediated the ubiquitination and degradation of YAP in a GSK-3beta-dependent manner. J Biol Chem. 2025;301(10):110652.

44. Sainero-Alcolado L, Liano-Pons J, Ruiz-Perez MV, and Arsenian-Henriksson M. Targeting mitochondrial metabolism for precision medicine in cancer. Cell Death Differ. 2022;29(7):1304–17.

45. Sousa B, Pereira J, and Paredes J. The Crosstalk Between Cell Adhesion and Cancer Metabolism. Int J Mol Sci. 2019;20(8).

46. Querzoli P, Coradini D, Pedriali M, Boracchi P, Ambrogi F, Raimondi E, et al. An immunohistochemically positive E-cadherin status is not always predictive for a good prognosis in human breast cancer. Br J Cancer. 2010;103(12):1835–9.

47. Burandt E, Lubbersmeyer F, Gorbokon N, Buscheck F, Luebke AM, Menz A, et al. E-Cadherin expression in human tumors: a tissue microarray study on 10,851 tumors. Biomark Res. 2021;9(1):44.

48. Possemato R, Marks KM, Shaul YD, Pacold ME, Kim D, Birsoy K, et al. Functional genomics reveal that the serine synthesis pathway is essential in breast cancer. Nature. 2011;476(7360):346–50.

49. Koo JH, and Guan KL. Interplay between YAP/TAZ and Metabolism. Cell Metab. 2018;28(2):196–206.

50. Kang S, Antoniewicz MR, and Hay N. Metabolic and transcriptomic reprogramming during contact inhibition-induced quiescence is mediated by YAP-dependent and YAP-independent mechanisms. Nat Commun. 2024;15(1):6777.

51. Kashihara T, Mukai R, Oka SI, Zhai P, Nakada Y, Yang Z, et al. YAP mediates compensatory cardiac hypertrophy through aerobic glycolysis in response to pressure overload. J Clin Invest. 2022;132(6).

52. Thrash HL, and Pendergast AM. Multi-Functional Regulation by YAP/TAZ Signaling Networks in Tumor Progression and Metastasis. Cancers (Basel). 2023;15(19).

53. Ghaboura N. Unraveling the Hippo pathway: YAP/TAZ as central players in cancer metastasis and drug resistance. EXCLI J. 2025;24:612–37.

54. Luo J, Deng L, Zou H, Guo Y, Tong T, Huang M, et al. New insights into the ambivalent role of YAP/TAZ in human cancers. J Exp Clin Cancer Res. 2023;42(1):130.

55. Zangari J, Petrelli F, Maillot B, and Martinou JC. The Multifaceted Pyruvate Metabolism: Role of the Mitochondrial Pyruvate Carrier. Biomolecules. 2020;10(7).

56. Kehrberg RJ, and DeMali KA. E-Cadherin: A conductor of cellular signaling. Curr Opin Cell Biol. 2025;95:102559.

57. Lin WH, Cooper LM, and Anastasiadis PZ. Cadherins and catenins in cancer: connecting cancer pathways and tumor microenvironment. Front Cell Dev Biol. 2023;11:1137013.

58. Benham-Pyle BW, Pruitt BL, and Nelson WJ. Cell adhesion. Mechanical strain induces E-cadherin-dependent Yap1 and beta-catenin activation to drive cell cycle entry. Science. 2015;348(6238):1024–7.

59. Arnaud T, Rodrigues-Lima F, Viguier M, and Deshayes F. Interplay between EGFR, E-cadherin, and PTP1B in epidermal homeostasis. Tissue Barriers. 2023;11(3):2104085.

60. Houtekamer RM, van der Net MC, Vliem MJ, Noordzij T, van Uden L, van Es RM, et al. E-cadherin mechanotransduction activates EGFR-ERK signaling in epithelial monolayers by inducing ADAM-mediated ligand shedding. Science signaling. 2025;18(886):eadr7926.

61. Qian X, Karpova T, Sheppard AM, McNally J, and Lowy DR. E-cadherin-mediated adhesion inhibits ligand-dependent activation of diverse receptor tyrosine kinases. EMBO J. 2004;23(8):1739–48.

62. Rubsam M, Mertz AF, Kubo A, Marg S, Jungst C, Goranci-Buzhala G, et al. E-cadherin integrates mechanotransduction and EGFR signaling to control junctional tissue polarization and tight junction positioning. Nat Commun. 2017;8(1):1250.

63. Zhao Y, Montminy T, Azad T, Lightbody E, Hao Y, SenGupta S, et al. PI3K Positively Regulates YAP and TAZ in Mammary Tumorigenesis Through Multiple Signaling Pathways. Mol Cancer Res. 2018;16(6):1046–58.

64. Gargalionis AN, Papavassiliou KA, Basdra EK, and Papavassiliou AG. Update on the relevance of mechanobiological mechanisms in lung cancer. Transl Oncol. 2025;55:102375.

65. Zhao Z, Mei Y, Wang Z, and He W. The Effect of Oxidative Phosphorylation on Cancer Drug Resistance. Cancers (Basel). 2022;15(1).

66. Cadassou O, and Jordheim LP. OXPHOS inhibitors, metabolism and targeted therapies in cancer. Biochem Pharmacol. 2023;211:115531.

67. Kalyanaraman B, Cheng G, Hardy M, and You M. OXPHOS-targeting drugs in oncology: new perspectives. Expert Opin Ther Targets. 2023;27(10):939–52.

68. Coleman MF, Cotul EK, Pfeil AJ, Devericks EN, Safdar MH, Monteiro M, et al. Hypoxia-mediated repression of pyruvate carboxylase drives immunosuppression. Breast Cancer Res. 2024;26(1):96.

69. Han MT, Pei H, Sun QQ, Wang CL, Li P, Xie YY, et al. ZY-444 inhibits the growth and metastasis of prostate cancer by targeting TNFAIP3 through TNF signaling pathway. Am J Cancer Res. 2023;13(4):1533–46.

70. Lisci M, Vericel F, Liu Y, Gallart-Ayala H, Ivanisevic J, Skinner OS, et al. Functional nutrient-genetic profiling reveals biotin and FBXW7 are essential to bypass glutamine addiction. Mol Cell. 2026.

71. Cheng T, Sudderth J, Yang C, Mullen AR, Jin ES, Mates JM, et al. Pyruvate carboxylase is required for glutamine-independent growth of tumor cells. Proc Natl Acad Sci U S A. 2011;108(21):8674–9.

72. Kota P, Terrell EM, Ritt DA, Insinna C, Westlake CJ, and Morrison DK. M-Ras/Shoc2 signaling modulates E-cadherin turnover and cell-cell adhesion during collective cell migration. Proc Natl Acad Sci U S A. 2019;116(9):3536–45.

73. Thonpho A, Rojvirat P, Jitrapakdee S, and MacDonald MJ. Characterization of the distal promoter of the human pyruvate carboxylase gene in pancreatic beta cells. PLoS One. 2013;8(1):e55139.

74. Pore M, Balamurugan K, Atkinson A, Breen D, Mallory P, Cardamone A, et al. Assessment of Imaging Mass Cytometry (IMC) as a Tool to Characterize Circulating Tumor Cells (CTCs) in Preclinical Mouse Models. bioRxiv. 2025.

75. Bates D, Mächler M, Bolker B, and Walker S. Fitting Linear Mixed-Effects Models Using lme4. Journal of Statistical Software. 2015;67(1):1–48.

76. Kuznetsova A, Brockhoff PB, and Christensen RHB. lmerTest Package: Tests in Linear Mixed Effects Models. Journal of Statistical Software. 2017;82(13):1–26.

