## Supplemental Figures for "Metabolic maintenance of breast cancer cells and metastases through E-cadherin/YAP–dependent pyruvate carboxylase expression"

**Figure S1**

A.

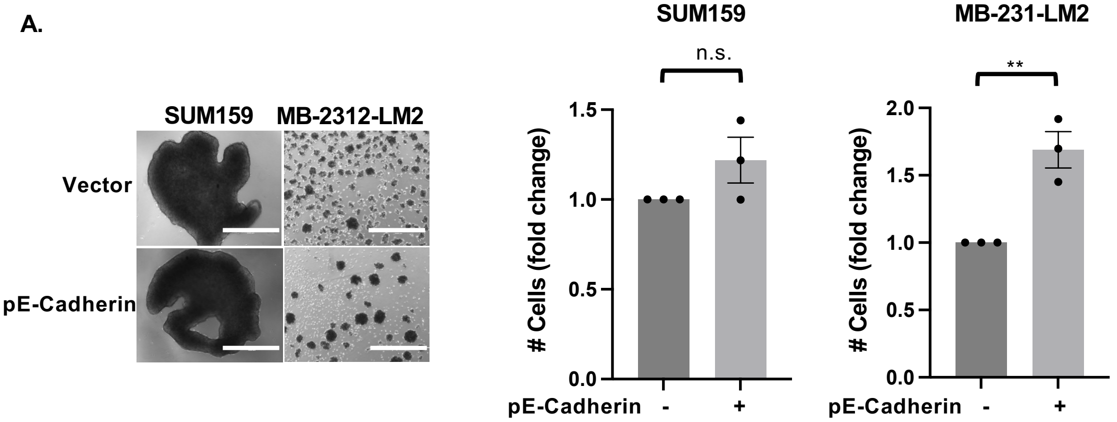


B.

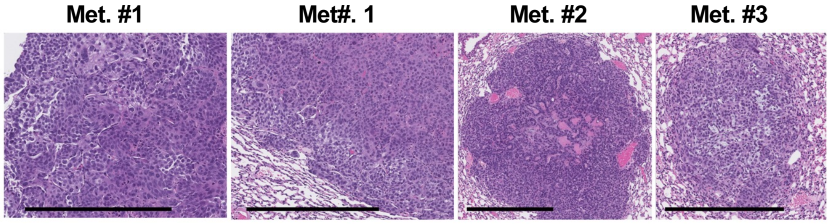


**Figure S1. E-Cadherin Promotes expression of pyruvate carboxylase**. **(A)** Light microscopy images of SUM159 and MDA-MB-231-LM2 cells after 3 days in EmC and transient transfection with E-cadherin expression construct or vector control (scale bar =1 mm) and quantification of viable cells from 3 independent experiments expressed as fold-change compared to vector control. Quantitative data are mean ± SEM, $**P<0.01$, n.s. non-significant. **(B)** Images of three H&E stained SUM149 experimental lung metastases (two regions representing metastasis #1) used for quantification of immunostaining (Figures 2F-G).

**Figure S2**

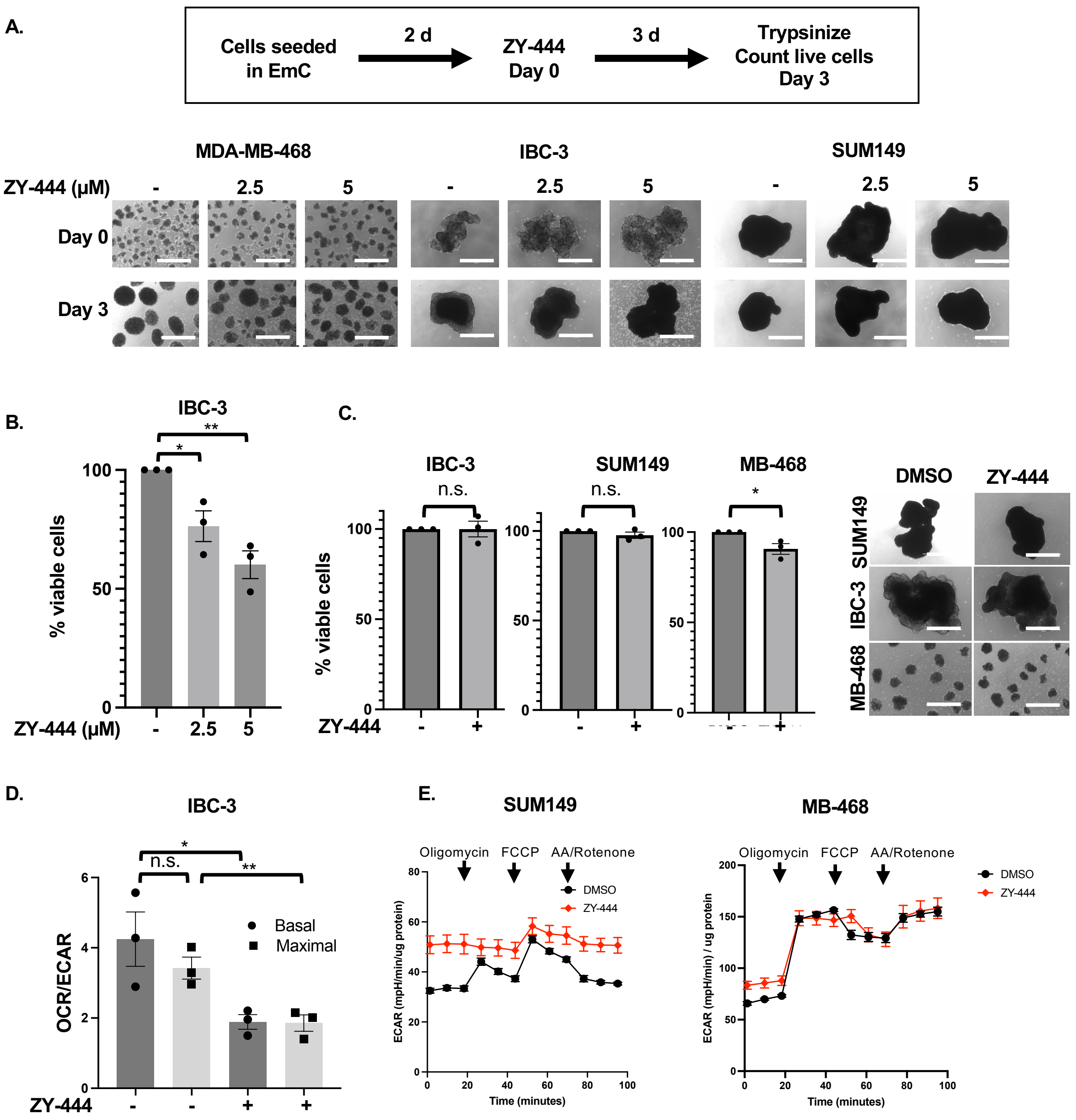


**Figure S2. Effect of the PC inhibitor ZY-444 on cell cultures.** **(A)** Schematic representation of experimental setup and light microscopy images of cells cultured in EmC before (Day 0) and after 3 days of exposure to ZY-444 (scale bar =1 mm). **(B)** Quantification of IBC-3 cells after three days of treatment with the indicated doses of ZY-444 compared to DMSO control. **(C)** Light microscopy images (right) and quantification (left) of the indicated cells after three days in EmC plus 12 h treatment with ZY-444 or DMSO control (scale bar= 1mm). **(D)** Basal and maximal OCR/ECAR ratios of IBC-3 cells as in panel A. **(E)** Representative Seahorse analysis of the extracellular acidification rate (ECAR) profiles from the experiment shown in Figure 3B. Quantitative data are mean ± SEM, **P*<0.05, ***P*<0.01, n.s. non-significant.

**Figure S3**

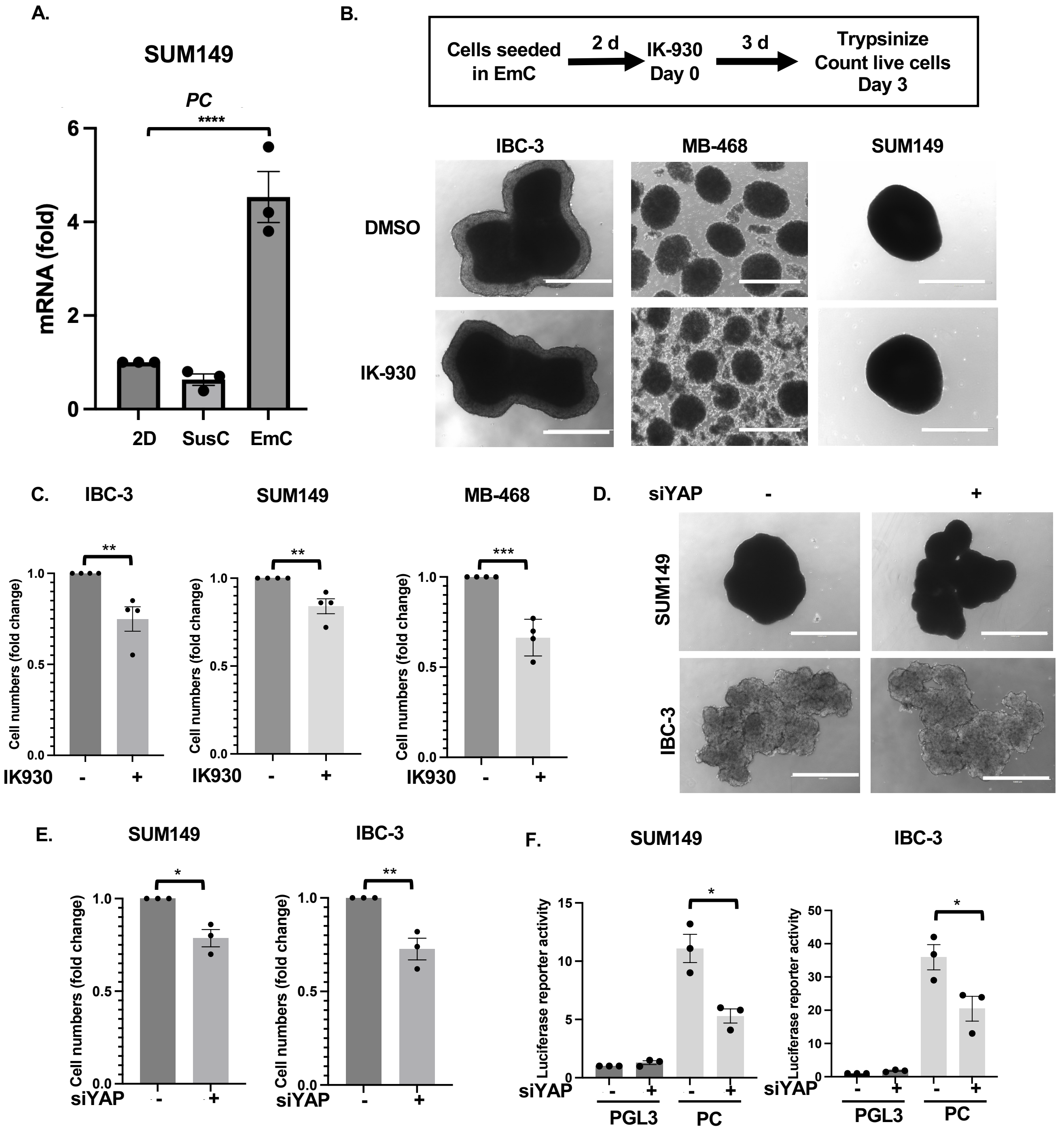


**Figure S3. The YAP/TEAD transcription factors support PC expression and OXPHOS.**
**(A)** qRT-PCR analysis of *PC* mRNA levels in SUM149 cells cultured in 2D, suspension (SusC) or EmC (n=3). **(B)** Schematic of experimental procedures and light microscopy images of EmC treated with IK-930 or DMSO control (scale bar = 1 mm). **(C)** Quantification of the indicated cells cultured as in panel B. **(D)** Light microscopy images of indicated cell lines transfected with control or siYAP after 3 days of EmC (scale bar =1 mm). **(E)** Quantification of SUM149 and IBC-3 cell cultures as in panel D. (**F**) Normalized luciferase reporter activities from the PC promoter ± siYAP in the indicated cell lines. Data are mean ± SEM, *$P$<0.05, **$P$<0.01, ***$P$<0.001, ****$P$<0.0001.

**Figure S4**

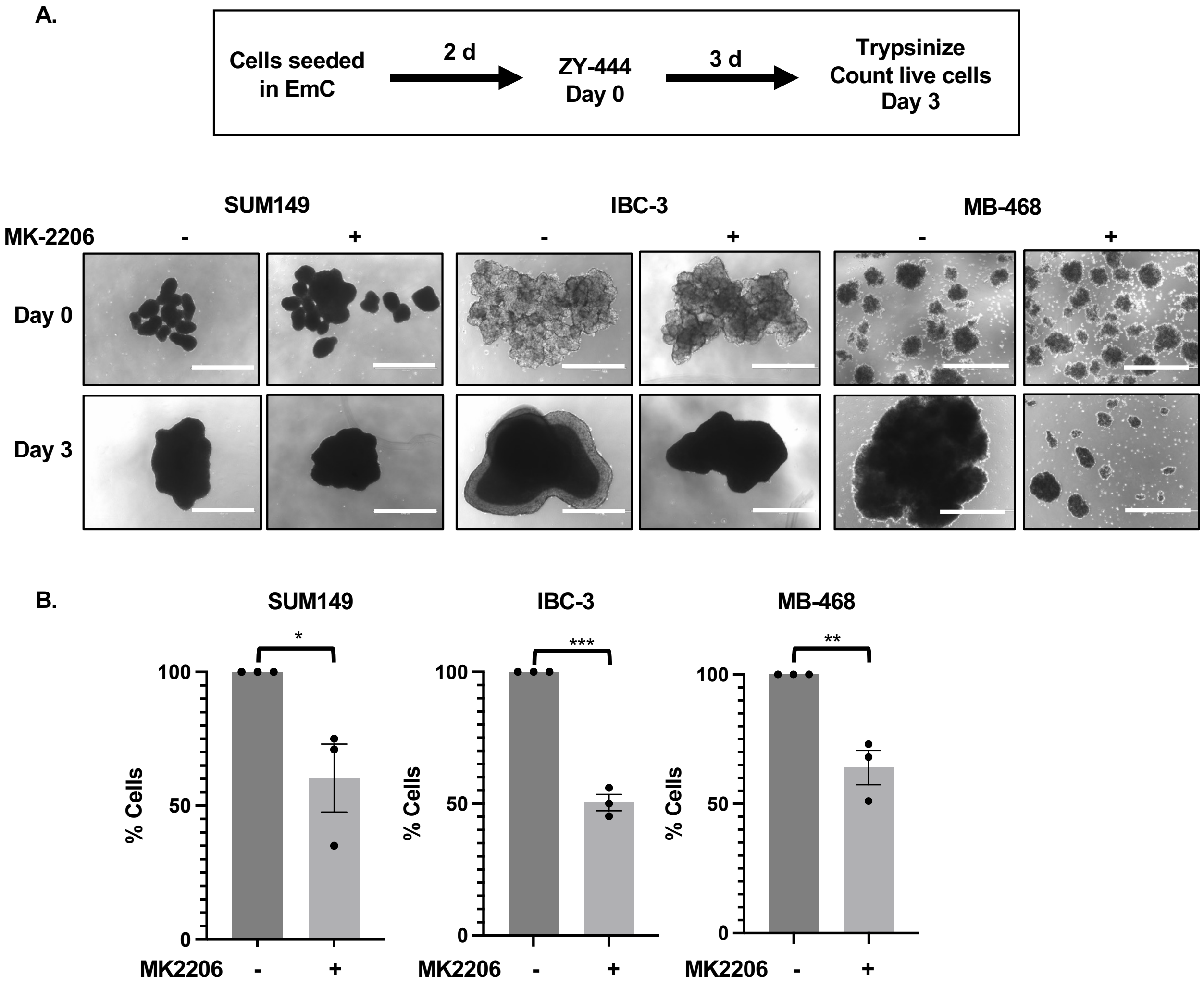


**Figure S4. Effect of MK-2206 on cells cultured in EmC (A)** Schematic of experimental procedures and light microscopy images of cells in EmC treated with MK-2206 or DMSO control (scale bar = 1 mm). **(B)** Quantification of cell cultures as in panel A. Data are mean ± SEM, **P*<0.05, ***P*<0.01, ****P*<0.001.

**Figure S5**

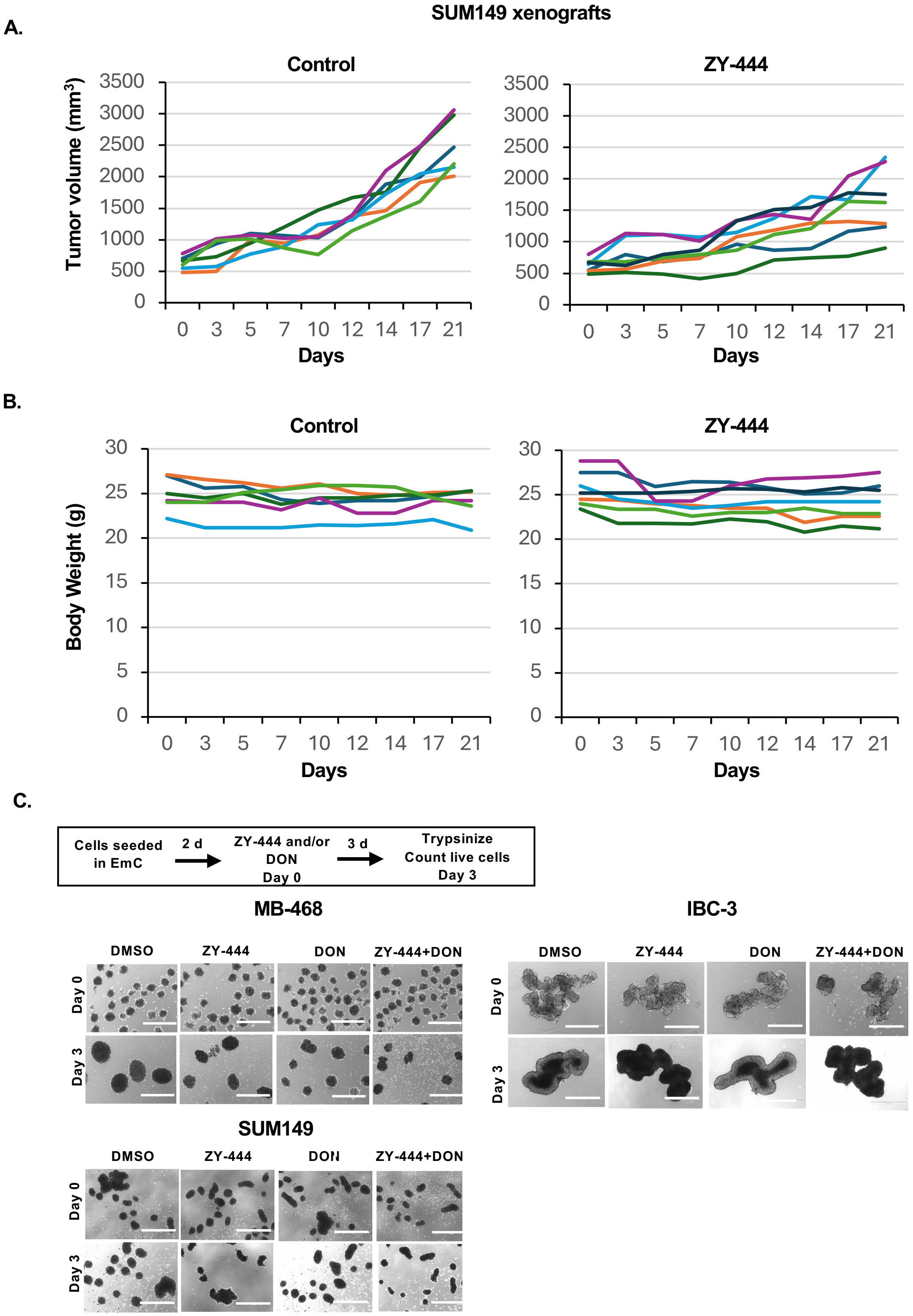


**Figure S5. Effect of ZY-444 in vivo and drug combinations in EmC. (A)** Tumor volume measurements of individual SUM149 xenografts in mice treated with DMSO (n=6) or ZY-444 (n=7) for 21 days. **(B)** Total body weights of mice as in panel A. **(C)** Schematic of experimental procedures and representative light microscopy images of EmC treated with ZY-444 (2.5 μM), DON (5 μM), both, or only DMSO as control (scale bar = 1 mm).
